# Constructing Virtual Viruses in A Spatial Minimal Cell

**DOI:** 10.64898/2026.09.22.753465

**Authors:** Weize Sun, Yong Tang, Mingyue Qiu, Jintao Li, Fei Luo, Ce Dou

## Abstract

Viral development links gene expression and genome replication to particle formation within a cell. Here we construct virtual viruses in a JCVI-syn3A-based spatial minimal cell. For mycoplasma virus P1, molecular events connect expression and replication to coarse-grained packaging, followed by a spatial reconstruction of membrane opening and particle release. Individual finite-size particles coexist, move across the boundary and undergo partial crossings and returns before complete exit. In contrast, Acholeplasma phage MV-L1 (L1) expresses its genes and accumulates genomes as the host continues to grow and develops a constricted shape. The coexistence of viral activity and continued host growth is qualitatively consistent with the non-lytic biology of group 1 acholeplasmaviruses. These reconstructions extend the virtual cell to the viruses that develop within it, linking molecular activity, particle behavior and host growth.

## INTRODUCTION

A virus must express its genes, copy its genome and organize its products into particles before progeny can leave the cell. These processes share a cellular environment, although their molecular and spatial requirements are often studied separately. Whole-cell models offer a way to connect molecular reactions with cellular behavior [1,2], providing a framework for studying viral production together with the host cell.

Genome minimization has made such cellular reconstructions more tractable [3,4]. Models of JCVI-syn3A connect metabolism and information processing to spatial organization [5], and the recent four-dimensional reconstruction follows these processes through growth and division [6]. Together with chromosome geometry and stochastic reaction–diffusion methods [7,8], this framework makes it possible to ask how an additional genetic program develops within a spatial cell, and how its products become particles that occupy and eventually leave that cell.

Intracellular phage models have connected viral production to host physiology and genome organization [9,10,11]. TABASCO and Pinetree resolve transcription and translation along a genome [12,13], and a recent ϕX174 reconstruction examined viral regulatory elements against transcript data [14]. However, connecting these molecular descriptions to particle formation and release within a growing spatial minimal cell requires a representation of both viral products and the changing cellular boundary. Such a representation would allow viral development and host growth to be studied together in a common spatial environment.

Here we present virtual viruses based on mycoplasma virus P1 [15] and Acholeplasma phage MV-L1 (L1) [16] within a JCVI-syn3A-based spatial minimal cell. The P1 reconstruction connects gene expression and genome replication to coarse-grained packaging, membrane opening and complete-particle release. By contrast, L1 continues to express its genes and accumulate genomes while the host grows and develops a constricted shape. The coexistence of L1 activity and continued host growth is qualitatively consistent with the non-lytic biology of group 1 acholeplasmaviruses [16,17]. Together, these virtual viruses extend minimal-cell reconstruction from the cell’s own life cycle to the viral processes it can support.

## RESULTS

### Establishing virtual viruses in a spatial minimal cell

The virtual viruses develop within an explicit spatial minimal cell. The initial configuration contains chromosome coordinates, ribosome centers and host proteins (Fig. 1A). Their radial organization and the abundance distribution of 455 host protein species describe the inherited cellular environment (Fig. 1C,D; Fig. S1). Viral annotations specify the added molecular program; packaging and placement events then connect that program to the spatial particle model (Fig. 1A,G).

**Figure 1.**
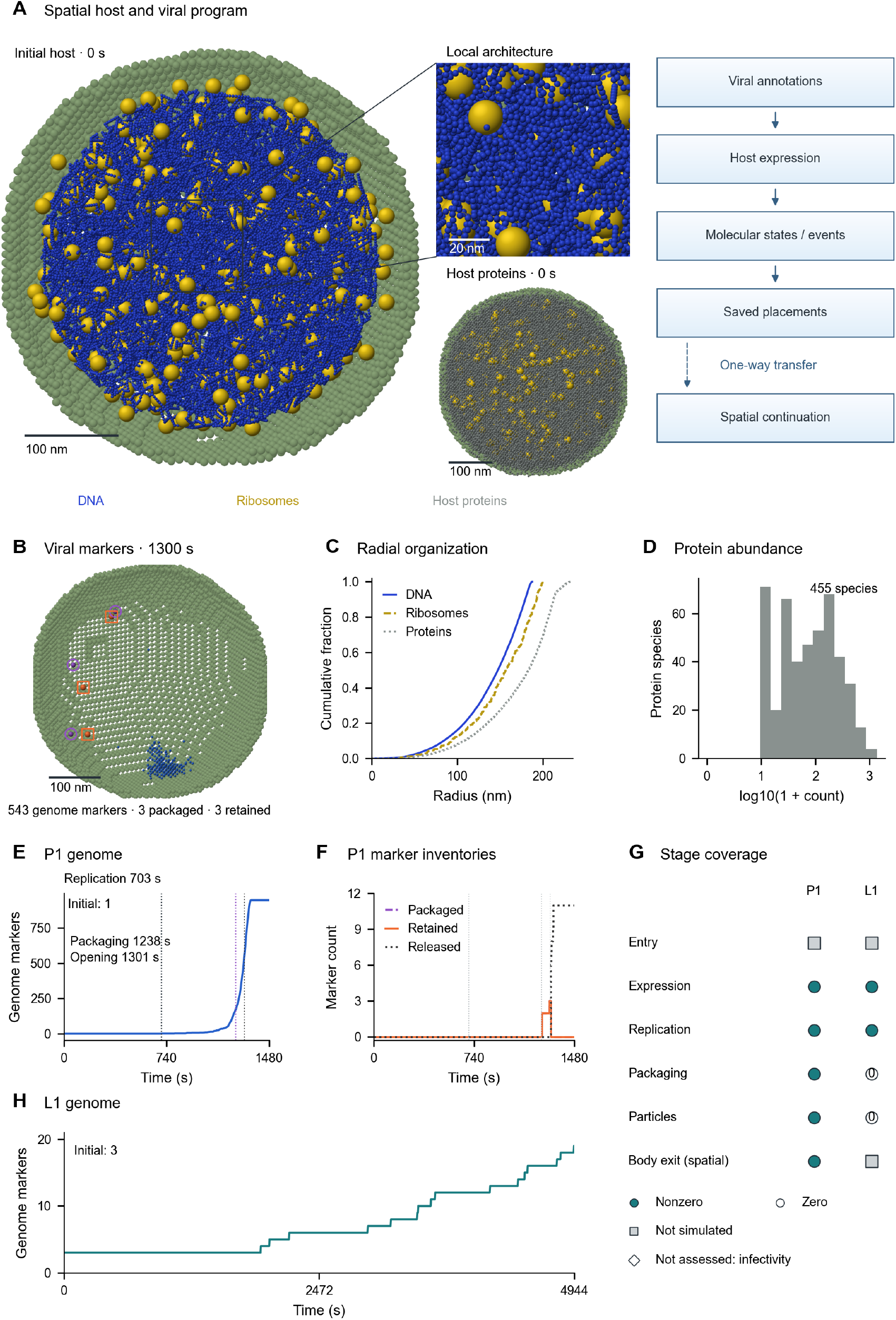
Virtual viruses within a spatial minimal cell. (A) Initial host geometry with a registered 100 × 100-nm local enlargement, a separate host-protein view and the viral-program interface. The dashed arrow denotes one-way transfer to the spatial continuation. (B) P1 markers at 1300 s: 543 genome markers, three packaged and three retained. Purple circles and orange squares locate packaged and retained coordinates, including projected overlaps. (C) Empirical cumulative radial distributions of all initial chromosome, ribosome and protein coordinates. (D) Abundances of 455 host protein species, displayed as log10(1 + count); translation-cost states are excluded. (E,F) P1 genome and downstream marker inventories over 0–1480 s. Labels identify replication at 703 s, packaging at 1238 s and opening at 1301 s. (G) Stage coverage distinguishes nonzero, zero, not simulated and not assessed. Body exit belongs to the P1 spatial continuation. (H) L1 genome-marker inventory over 0–4944 s; downstream inventories remain zero (Fig. S4D). Initial genomes are one for P1 and three for L1. Spatial cutaways affect display only; scale bars denote nanometers.

In the P1 trajectory, gene expression precedes genome amplification and packaging (Fig. 1E,F). Transcription and translation first occur at 13 and 25 s, replication at 703 s and packaging at 1238 s (Table S1). Eleven packaging events have corresponding placements over the 0–1480-s record. The terminal inventories contain 947 genome markers and eleven released markers (Fig. 1E,F). The spatial continuation follows the resulting particles beyond these molecular readouts (Fig. 1G).

L1 expresses its four loci and accumulates genomes from 3 to 19 over 0–4944 s, while the three downstream particle inventories remain zero (Fig. 1H; Fig. S4D). Its first replication event occurs at 1907 s, beyond the shared 0–1480-s window displayed in Fig. S6C. These distinct trajectories provide a particle-producing P1 case and an L1 case in which molecular activity continues during host growth (Fig. 4H–J).

### From viral genes to transcripts and proteins

The viral annotations specify the loci followed through synthesis and molecular accumulation. The eleven P1 loci and four L1 loci retain their genomic coordinates, including the two origin-spanning segments of L1_4 (Fig. 2A). Each locus defines gene, transcript and protein states; sequence length enters completion rates, and sequence composition specifies resource-cost requests. The P1 configuration also applies translation weights to selected loci (Fig. 2B; Tables S6,S7,S9).

**Figure 2.**
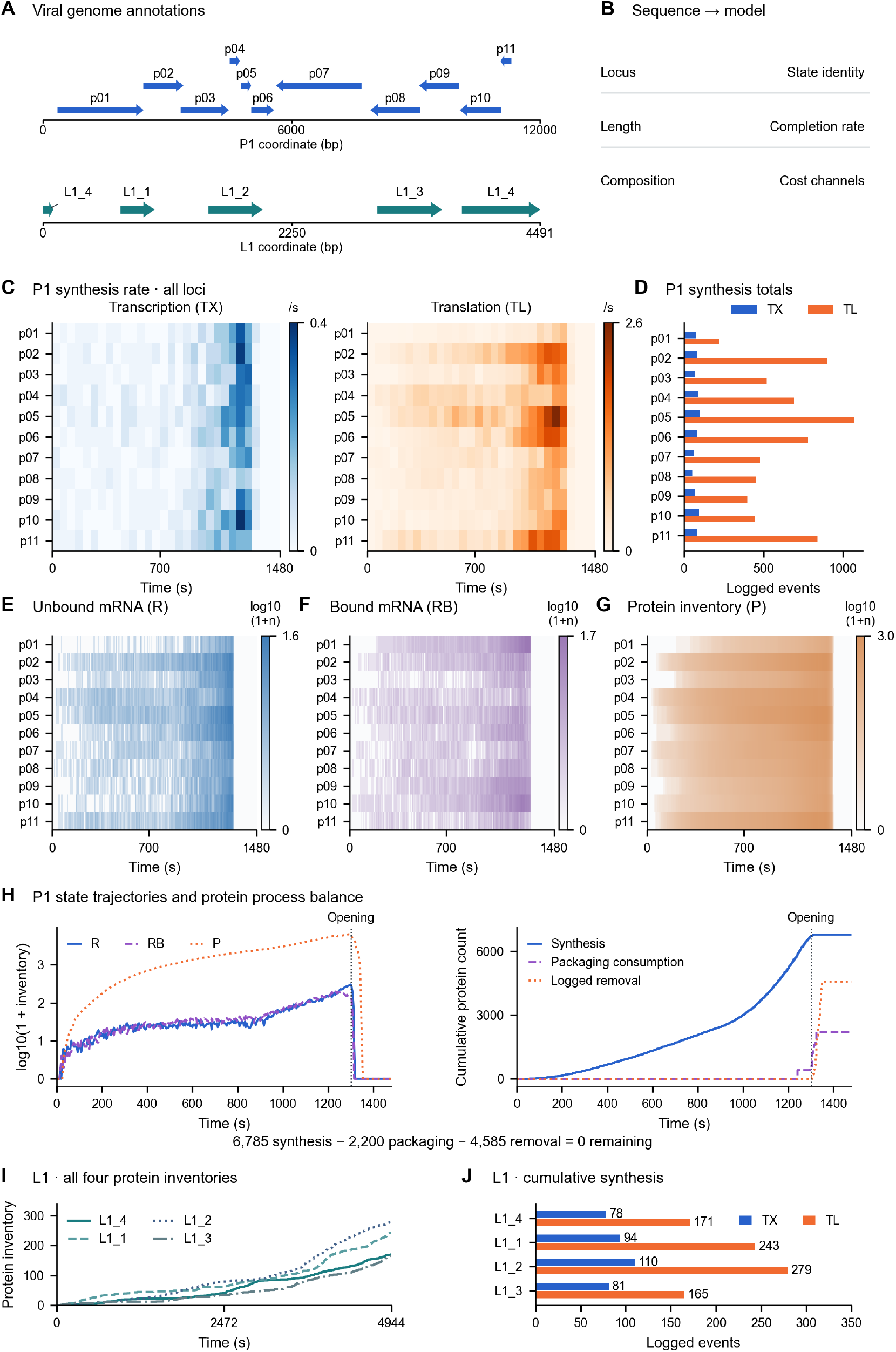
Gene-resolved expression in P1 and L1. (A) Coordinate tracks for eleven P1 loci and four L1 loci. Both origin-spanning L1_4 segments identify the same gene. (B) Sequence-to-model mapping in the source implementation described in Methods; state, rate and cost definitions are provided in Tables S6 and S7. (C) Gene-resolved transcription (TX) and translation (TL) rates. Event multiplicities are summed in left-closed, right-open 50-s bins and divided by their actual widths; the final 1450–1480-s bin spans 30 s and includes the terminal edge. Independent rate scales are shown. (D) Cumulative TX and TL totals. (E–G) Unbound mRNA (R), ribosome-bound mRNA (RB) and protein (P) inventories, including zeros, on independent log10(1 + count) scales without row normalization. (H) P1 expression and protein fates: aggregate state trajectories and cumulative synthesis, packaging consumption and removal over 0–1480 s; dotted lines mark opening at 1301 s. Of 6,785 synthesized proteins, 2,200 are consumed during packaging and 4,585 undergo logged removal, leaving zero inventory. Consumption is a configured 200 subunits per packaging event. (I,J) L1 protein trajectories and per-locus synthesis totals across the complete 0–4944-s window. TX/TL colors match D. Complete L1 R/RB/P trajectories are provided in Fig. S4.

P1 synthesis is uneven across loci (Fig. 2C,D). MpP1p05 contributes 1,070 of the 6,785 translation events, whereas MpP1p01 contributes 219; transcription totals range from 52 to 100 per locus, summing to 858 (Table S9). Thus, the spread in cumulative protein synthesis is larger than that in transcription. The locus annotation identifies MpP1p01 as DNA polymerase; the other product labels and the configured packaging requirements are listed separately from their translation weights (Table S9).

Unbound mRNA (R) and ribosome-bound mRNA (RB) vary through time, while proteins accumulate before the decline associated with packaging and the opening rule (Fig. 2E–H; Fig. S2). Aggregate R, RB and P inventories peak at 304, 196 and 6,236, respectively (Fig. 2H). RNAP and ribosome binding and release couple expression to the host machinery (Fig. S6D). Protein synthesis and protein inventory diverge because packaging consumes part of the synthesized pool and the post-opening removal rule removes the remainder: 2,200 proteins enter configured packaging and 4,585 undergo logged removal, leaving zero endpoint inventory (Fig. 2H; Fig. S6B). The packaging rule consumes 200 structural subunits per event.

L1 produces 363 transcription and 858 translation events across its four loci (Fig. 2I,J). Its protein inventories accumulate with synthesis, alongside the changing transcript states shown in Fig. S4E–H. Gene expression and genome accumulation therefore continue through a window with no recorded particle output (Fig. 1H; Fig. S4D; Table S4). Video S2 places this molecular activity in the growing host geometry.

### Linking molecular events to spatial particles

To follow viral development beyond molecular production, we linked packaging and placement events to individual particles in the spatial cell. In the P1 reconstruction, genome, packaged and retained markers provide parallel molecular inventories (Fig. 3A). The accompanying placement records supply each particle’s identity, position and orientation, together with its scheduled placement time. The spatial continuation constructs the particle and admits it when the geometric rules permit, preserving its identity if admission is deferred (Fig. 3B).

**Figure 3.**
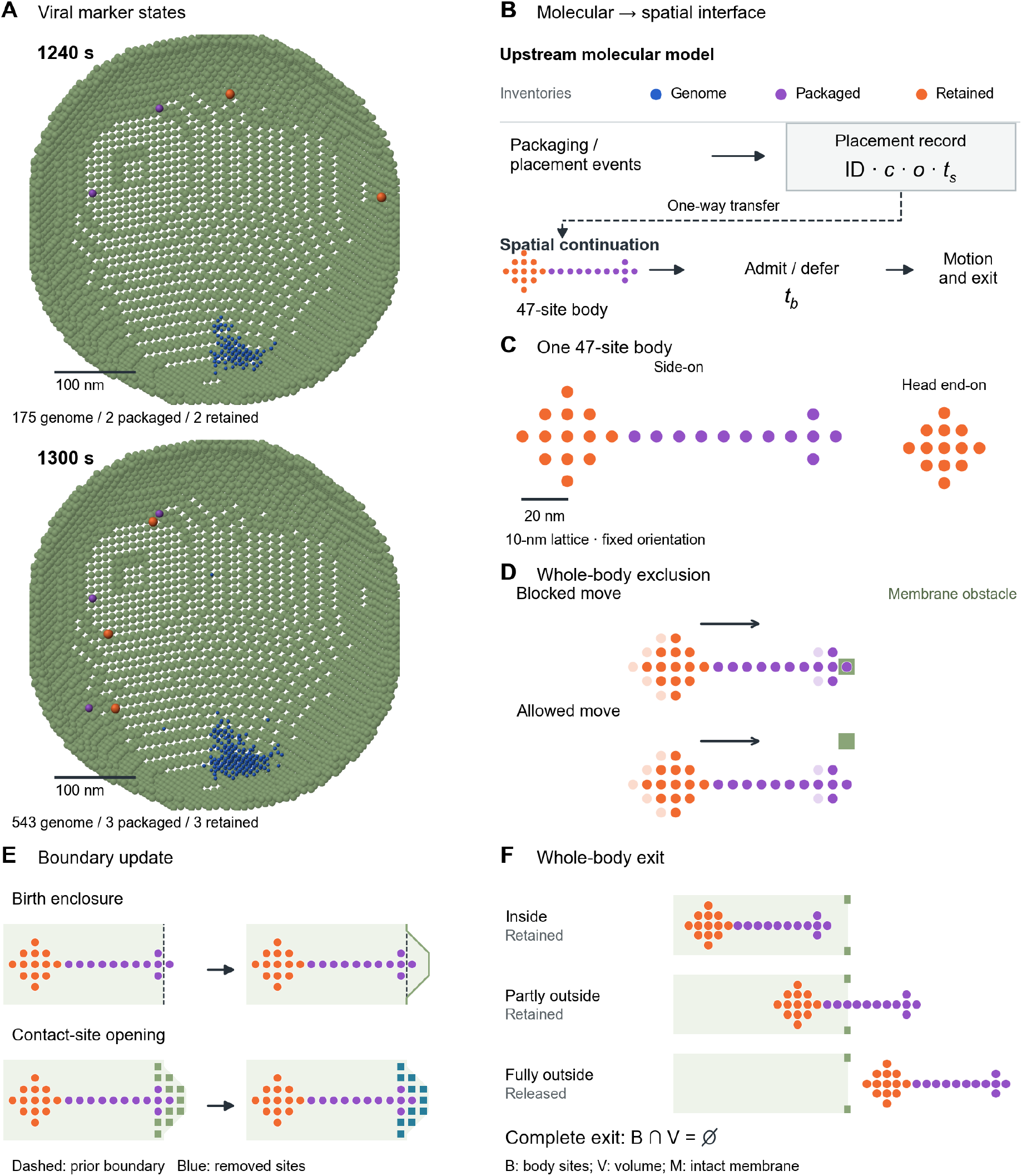
From molecular events to spatially resolved particles. (A) P1 marker distributions at 1240 and 1300 s, with a common camera, coordinate transformation and display cutaway; counts give genome, packaged and retained inventories. Scale bars, 100 nm. (B) Packaging and placement events supply identity, position *c*, orientation *o* and scheduled time *t*_s_ to construct each spatial body. The continuation admits or defers the body and determines accepted birth time *t*_b_, preserving its identity. Genome, packaged and retained inventories are parallel readouts. The dashed connection denotes one-way transfer. (C) Side and head-end projections of the same 47-site body on a 10-nm lattice, with fixed orientation; projected sites can overlap. Sites represent occupied volume, not the 200 protein subunits consumed per packaging event. (D) A one-site translation proposal is blocked by or clears a membrane obstacle. Pale bodies show prior positions and solid bodies show proposals; arrows indicate direction and are not displacement scale bars. (E) Before–after rule examples of birth enclosure and contact-site opening. Dashed and solid contours identify prior and updated boundaries; green sites remain intact and blue sites are removed. (F) Inside and partly external bodies remain retained; complete exit requires every occupied site to lie outside the current volume, equivalently B ∩V is empty. B denotes body sites, V cell volume and M intact membrane. Orange and purple encode head and tail/baseplate structure. C–F show model geometry and rule schematics; the event-resolved trajectory is shown in Fig. 4D.

Each particle has a head, tail and baseplate represented by 47 occupied sites on a 10-nm lattice (Fig. 3C). This finite-size representation gives the particle an explicit shape and occupied volume, distinct from the 200 protein subunits consumed per packaging event. Movement is permitted only when the whole particle clears membrane obstacles (Fig. 3D). Admission updates the enclosing cell boundary; after the prescribed opening gate, membrane sites in accumulated contact neighborhoods are removed, changing the routes available for particle motion (Fig. 3E).

Tracking the whole particle also distinguishes partial crossing from complete exit. A particle remains retained while any occupied site lies within the current cell volume and is classified as released only when all sites lie outside (Fig. 3F). Molecular production thus supplies identifiable spatial particles whose movement and complete exit can be followed through time. The resulting trajectories are examined in Fig. 4, while upstream marker distributions and distances are reported separately in Fig. S5.

**Figure 4.**
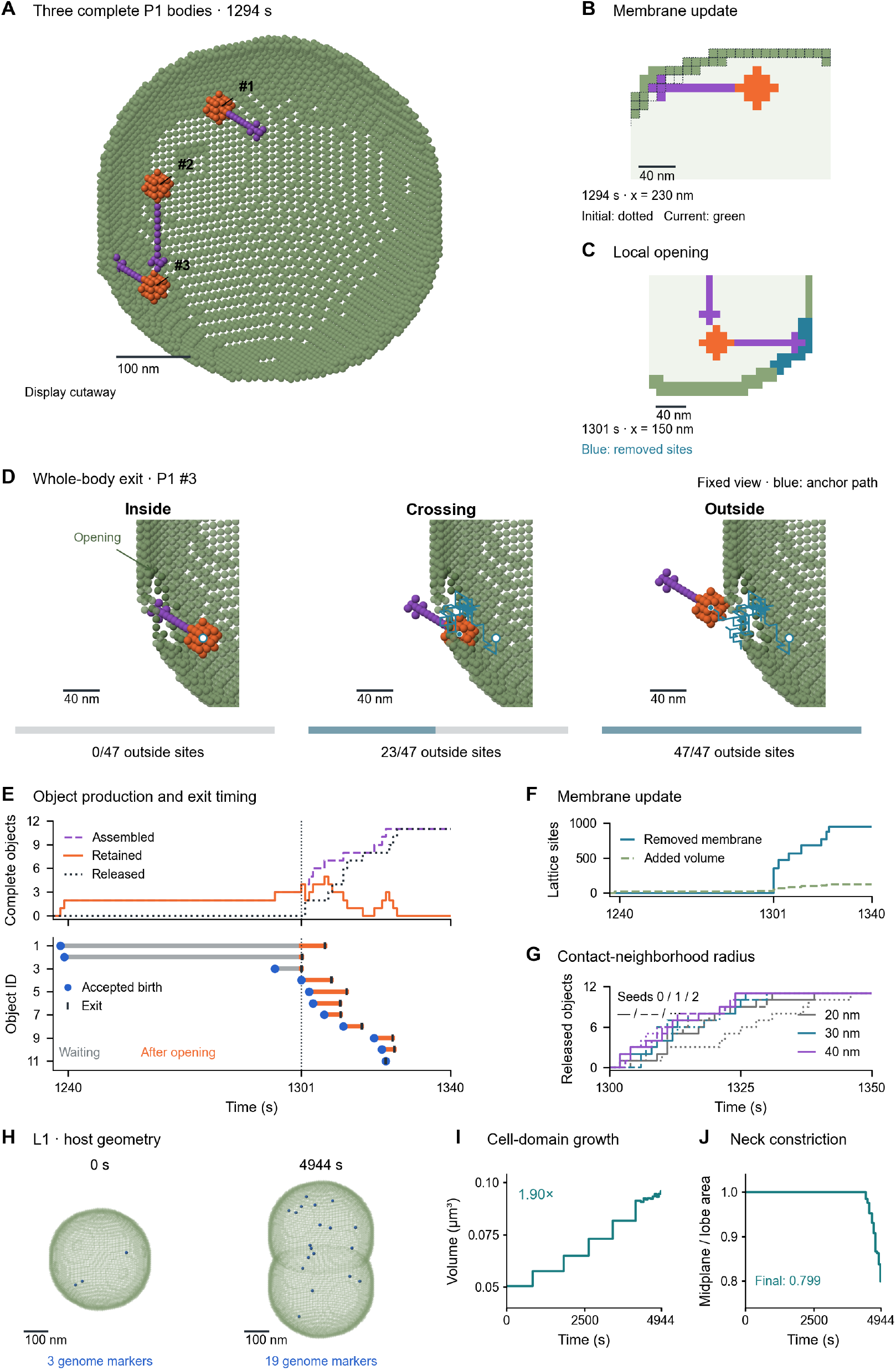
P1 particle release and L1 activity during host growth. (A) Three P1 bodies at 1294 s in the main continuation (30-nm contact neighborhood, seed 0), with a membrane display cutaway. (B,C) Membrane enclosure and opening sections; dotted outlines mark the initial boundary and blue sites mark removed membrane. (D) Event-resolved exit of particle #3, shown inside, crossing and outside in a fixed view with a membrane display cutaway. The blue line traces recorded head-anchor positions; the open circle marks the starting position. Bars report sites outside the three-dimensional cell volume (0/47, 23/47 and 47/47). Orange and purple denote head and tail/baseplate sites. States are ordered in time but not equally spaced. (E) Complete-object counts and all 11 accepted-birth–exit histories. Gray denotes waiting for opening, orange post-opening transport, blue circles accepted birth and ticks exit; assembled = retained + released. (F) Removed membrane and added volume sites. (G) Nine continuations sharing upstream events: color identifies contact-neighborhood radius and line style identifies seed. (H) Historical L1 host geometry at 0 and 4944 s, shown at the same scale without a cutaway; blue marks genomes. (I) Cell-domain volume, including membrane sites. (J) Axial midplane area divided by the mean maximum cross-sectional area of the two axial halves. H–J describe one L1 trajectory; I,J show all 1237 checkpoints without smoothing. Spatial bars denote nanometers; object-level summaries and L1 measurements are provided in Fig. S7 and Table S4.

### P1 particle release and L1 activity during host growth

The P1 spatial continuation resolves body coexistence and exit against an evolving boundary. Starting from the parent geometry at 1236 s, it accepts the 11 saved placement events and contains three complete bodies at 1294 s (Fig. 4A). Accepted births update the enclosing boundary (Fig. 4B). The prescribed gate opens contact neighborhoods at 1301 s and simultaneously enables stochastic translation; bodies remain fixed before this gate (Fig. 4C). The main condition uses a 30-nm contact neighborhood and spatial seed 0.

Event-resolved replay follows object #3 from wholly inside, through a partly external configuration, to complete exit (Fig. 4D). The three displayed states contain 0, 23 and 47 external sites at 0, 25 and 32 ms after opening. All 11 bodies exit by 1325.278 s in the main condition (Fig. 4E). Birth enclosure adds 126 volume sites and the final opening comprises 950 removed membrane sites (Fig. 4F). Video S1 presents the P1 molecular record and the separately computed conditional assembly and release segments.

Object histories separate waiting for the prescribed gate from transport after opening (Fig. 4E; Fig. S7). Measured from the later of accepted birth and gate opening to complete exit, the main-condition median interval is 4.824 s; condition medians span 1.109–8.579 s across the nine radius–seed combinations (Fig. 4G; Fig. S7D). All 11 objects per condition are included, including admission delayed until after opening. These intervals separate post-opening transport from pre-opening waiting under the imposed rules.

Crossing the boundary was not always a one-way event. The occupied head anchor returned inside before complete exit in 80 of the 99 condition–object records (Table S8). As a result, cumulative first-crossing counts exceeded complete-exit counts by up to four objects during the trajectories. Both readouts reached 11 objects per condition by 1480 s. Complete-body exit followed first anchor crossing by a median of 24.850 ms across all records (range, 0.032 ms– 28.106 s). The paired readouts resolve partial crossing and return on the same simulated paths; the full timing summaries are provided in Table S8.

L1 gene expression and genome accumulation coexist with continued host growth and division-associated constriction over the recorded interval (Fig. 4H–J; Video S2). Cell-domain volume, including membrane sites, increases from 0.050397 to 0.095845 μm^3^, or 1.90-fold. Over the same interval, the genome-marker inventory increases from 3 to 19 (Fig. 1H), giving a 3.33-fold rise in markers per unit cell-domain volume (Table S4). Genome accumulation therefore exceeds the proportional increase in host volume. The historical log records 10 division-associated geometry updates beginning at 4288.25 s, with division_started true at the endpoint (Table S4). The axial midplane-to-lobe area ratio decreases to 0.799 (Fig. 4J). Thus, the host enters division-associated shape updating during ongoing L1 activity, although the saved cell domain remains connected through 4944 s and daughter-cell separation is not observed.

## DISCUSSION

Constructing virtual viruses extends minimal-cell reconstruction to viral development within the cell. In the P1 reconstruction, gene expression and genome replication supply a molecular record that is connected through coarse-grained packaging and placement to finite-size particles. Following these particles distinguishes molecular production from spatial release: protein synthesis does not by itself specify particle output, and an initial boundary crossing does not establish complete exit (Figs. 2H, 3B and 4D). Individual bodies can cross partially and return before leaving the cell. This connection makes the intervening steps explicit, providing a framework for examining how molecular activity becomes particle output under specified cellular and boundary conditions.

L1 illustrates a complementary course in which viral expression and genome accumulation coexist with continued host growth and entry into division-associated shape updating (Fig. 4H–J; Table S4). This coexistence is qualitatively consistent with the non-lytic biology of group 1 acholeplasmaviruses, whose progeny can be released while host integrity is maintained [16,17]. No L1 particle output was recorded under the archived configuration; this trajectory is therefore used to examine viral activity alongside host growth rather than to infer L1 particle production or release. Membrane-associated DNA intermediates described for MVL51 provide biological context for examining the connection between replication and particle production [18].

Earlier intracellular T7 models connected genome organization to the timing of molecular synthesis and progeny production [10,11]. Incorporating host physiological parameters further showed how changes in cellular resources could alter phage growth [9]. These studies established that viral output depends on both the genetic program and the host conditions supporting it, and they compared model predictions with measured phage behavior. VirtualVirus builds on this line of inquiry by placing viral activity within an explicit minimal-cell environment and following the spatial fate of its particle products. The added representation resolves object occupancy and boundary transit, extending existing molecular models toward spatial particle behavior.

Gene-expression simulators address another part of this problem at finer molecular resolution. TABASCO follows individual DNA-associated molecules and base-pair-resolved transcription, while Pinetree resolves stochastic transcription and translation with individual polymerases and ribosomes and codon-specific translation rates [12,13]. The recent ϕX174 model used transcript measurements to fit regulatory parameters and compare alternative descriptions of genome regulation [14]. These approaches provide detailed links between genome organization and expression kinetics. Here, annotated loci define transcript and protein states, completion rates and resource requests within the inherited host model, and the resulting molecular events feed the particle representation. Our emphasis is therefore on connecting expression to subsequent spatial processes. The P1 translation weights and packaging requirements provide a coarse-grained link between expression and particle formation rather than a fitted regulatory model.

Spatial whole-cell reconstructions provide the cellular context for these viral processes. The JCVI-syn3A models integrate metabolism and information processing with spatial organization; the 4D reconstruction further follows host growth, chromosome dynamics and division [5,6]. Those host capabilities are inherited here. The added viral programs and the P1 continuation extend the description to packaging-linked object admission, boundary modification and complete-particle exit. A related spatial reaction–diffusion model of HBV infection examines intracellular viral chemistry and drug interactions in a reconstructed hepatocyte [19]. The relevant distinction is the set of processes represented and coupled, rather than the whole-cell label alone. In our P1 construction, molecular placements drive a one-way spatial continuation: the opening and mobility gate is prescribed, and subsequent particle motion and boundary changes do not feed back into the upstream reaction model. This coupling generates explicit particle histories and provides a foundation for future bidirectional models of infection, host response and release.

A central question is when increased molecular production leads to more released particles, and when packaging, spatial admission or boundary access becomes limiting. T7 models have shown how host physiology can shape viral production [9], while studies of gene-expression burden and resource allocation provide ways to examine competition for cellular capacity [20,21,22]. These findings motivate matched computational comparisons in which molecular supply, packaging requirements or boundary conditions are varied while the remaining configuration is held fixed. Such comparisons could distinguish a limitation in synthesis from one in particle formation or transport. The current separation of molecular and spatial readouts defines quantities for those comparisons; the difference between genome and particle counts alone does not identify a limiting mechanism.

Virtual virus construction should extend beyond P1 and L1. Other viruses could be explored by adapting their gene programs, replication strategies, particle structures and routes of release to an appropriate cellular environment. Related phages offer a direct starting point, whereas more distant viruses would also require different host reconstructions. Such extensions would allow changes in viral genes or life-cycle steps to be examined alongside their consequences for particle production and host behavior. The aim is to connect a virus’s genetic program with how it develops in, and changes, the cell that supports it.

### Limitations of the study

The current reconstructions simplify particle motion and membrane remodeling and sample a limited set of viral trajectories. The archived upstream trajectory is interpreted as a process-level reconstruction rather than a calibrated prediction of viral yield. Future development should connect L1 replication to particle production and establish bidirectional coupling between viral activity, particle behavior and host-cell dynamics.

## Supporting information

P1 VirtualVirus Visualization

L1 VirtualVirus Visualization

## DATA AND MATERIALS AVAILABILITY

### Study materials

This computational study did not generate biological materials or reagents.

### Data and code availability

Research code, selected P1/L1 reference tables, and the geometry and placement inputs for the P1 spatial continuation are publicly available at https://github.com/dmaskhhh/VirtualVirus. The repository supports rerunning the spatial example and checking the included reference results; its README and docs/reproduction.md describe the available files and reproduction scope. The complete historical upstream archives, full figure-rendering records and video production package remain locally retained and are not included in this public release. Run provenance and calculation definitions are documented in Methods and the Supplemental Information [23,24].

## ACKNOWLEDGMENTS

This work was supported by the National Natural Science Foundation of China (82572785) and the NSFC Key Projects of the Regional Innovation and Development Joint Fund (U23A20413).

## AUTHOR CONTRIBUTIONS

Weize Sun: Methodology, software, formal analysis, visualization, and writing – original draft. Yong Tang: Methodology, data curation, and validation. Mingyue Qiu: Data curation, formal analysis, and validation. Jintao Li: Methodology and writing – review & editing. Fei Luo: Supervision and writing – review & editing. Ce Dou: Conceptualization, supervision, funding acquisition, project administration, and writing – review & editing.

## DECLARATION OF AI USE

AI-assisted tools supported manuscript drafting and analysis-script development. The authors are responsible for reviewing the text, code, references and interpretations and for the submitted content.

## SUPPLEMENTAL INFORMATION

Document S1. Figures S1–S7, Tables S1–S9, and the legends for Videos S1 and S2. Video S1. P1 genome replication, conditional assembly and complete-body release. Video S2. Historical L1 molecular expression and genome accumulation over 0–4944 s.

## METHODS

### Data, software and computational resources

| RESOURCE | SOURCE | IDENTIFIER |
| --- | --- | --- |
| Deposited data |  |  |
| Mycoplasma virus P1 reference genome | NCBI RefSeq; Tu et al. | NC_002515.1; eleven loci |
| Parent simulation record | This study | r12ce_g1_2500s; selected tables: GitHub inputs/reference; full historical archive not publicly deposited |
| Acholeplasma phage MV-L1 annotation | Archived RefSeq input | NC_001341.1; four loci; natural host: *Acholeplasma laidlawii* |
| Historical L1 record | This study | See Table S4; selected tables: GitHub inputs/reference; full historical archive not publicly deposited |
| Conditional spatial continuations | This study | r2_s0 to r4_s2; nine-condition summaries: GitHub inputs/reference; runnable example: src/spatial |
| Figure data and event ledgers | This study | Selected tables and event ledgers: GitHub inputs/reference; complete figure records retained locally |
| Software and algorithms |  |  |
| Minimal_Cell_4DWCM | Thornburg et al. (2026) | doi:10.1016/j.cell.2026.02.009; historical executable identity not bound |
| Lattice Microbes | Roberts et al. (2013) | doi:10.1002/jcc.23130 |
| Current analysis environment | Recorded environment | Python 3.11; NumPy 2.4.6; SciPy 1.17.1; Matplotlib 3.10.9 |
| Spatial kernel and data readers | Recorded environment | Numba 0.67.0; h5py 3.16.0; Pillow 12.2.0 |
| Membrane continuation and verification scripts | This study | GitHub src/spatial, analysis and tests |
| Other |  |  |
| Statistical scope | This study | One P1 and one historical L1 trajectory; nine P1 conditional continuations |

### Computational model and simulation scope

This study comprises computational biological modeling and visualization only. No wet-lab experiments, human participants or human-derived data were involved. The inherited host framework is a spatial JCVI-syn3A-based whole-cell model [5,6,7,8]. The mycoplasma virus P1 program is defined from accession NC_002515.1 and the associated genome report [15,25]. P1 initial conditions represent post-entry placement of one genome. Neither natural infection of JCVI-syn3A nor host compatibility is assumed from the annotation.

### Parent record and event extraction

The parent archive r12ce_g1_2500s contains 371 count and spatial checkpoints through 1480 s, a stored stop time of 1481 s and opening at 1301 s. The analyzed horizon is the recorded interval; the requested duration was 2500 s. Saved descriptors include initialGenomes = 1, placementSeed = 11012, dnaRngSeed = 42, assemblyRateOverride = 0.02 and structuralSubunitTotal = 200. Table S2 and the full configuration retain the other placement, packaging and permeability settings.

Counts were read from transposed counts_and_fluxes.csv; events came from gene, topology-replication, topology-packaging and physical-placement logs. Transcription and translation totals sum logged event values. Packaging and placement were aligned by time as corresponding operations. Genome inventory equals initial genomes plus cumulative replication minus packaging; packaged inventory equals cumulative packaging minus the parent released-marker readout; retained inventory equals cumulative placement minus that readout. These are parallel inventory balances. The first nonzero parent released readout falls between the 1300- and 1304-s samples.

Two implementation issues were identified in the archived upstream trajectory: free-count synchronization could alter selected host propensities, and template registry consumption was not always accompanied by removal of the corresponding active template. Because the archived trajectory was not regenerated, the quantitative influence of these behaviors was not determined. The present study therefore interprets this record as a process-level reconstruction rather than a calibrated prediction of viral yield. Technical details are retained under Upstream numerical interventions in the Supplemental Information.

### Historical L1 record extraction and provenance

The historical archive p7_l1_continuous7200_genome3_dmg50_05161111 specifies four loci, accession NC_001341.1, initialGenomes = 3, placementSeed = 70701 and dnaRngSeed = 42. Its 1237 samples at 4-s intervals end at 4944 s, despite the requested 7200-s duration. All 36 V_L1_ state rows were retained. L1_4 spans the circular origin and is displayed as two segments of one locus (Fig. S4A,E–H). The natural host is *Acholeplasma laidlawii* [16].

Transcription, translation and replication were quantified by summing saved gene-event values. Count tables were cross-checked against HDF5 SpeciesCounts using the saved species-index mapping, and cumulative replication and translation were compared with the corresponding genome and protein inventories. All three downstream marker rows were present and zero.

Configuration, annotation and event records are retained with the source tables (Fig. S4; Table S4).

The retained metadata specify packaging_genome_start = 50, packaging_unpacked_reserve = 25 and packaging_conversion_per_second = 5, and describe assembly as consuming a packaged genome and one copy of each required protein. These values are retained as archived configuration parameters and are not treated as biologically calibrated thresholds. A separate 60-s current-source feasibility run checks initialization and count consistency only, as documented in the supplemental computational record.

Host morphology was quantified from all 1237 saved L1 Sites arrays, spanning 16 cell/membrane geometries (Fig. 4H–J). The cell-domain mask includes every non-extracellular site, including membrane; its voxel count was multiplied by (10 nm)^3^ to obtain volume. Cross-sectional areas were measured along the stored z axis. The midplane is the rounded midpoint of the axial bounding box; the neck-area ratio is its area divided by the mean maximum area of the two axial halves. Genome-marker abundance per unit cell-domain volume is the recorded V_L1_Genome inventory divided by this volume at each shared timestamp, with fold changes relative to the initial value. Six- and 26-neighbor connectivity were evaluated for every geometry. The first division-associated log entry at 4288.25 s and the terminal division_started flag identify entry into shape updating; repeated DIVISION STEP COMPLETE entries denote geometry updates rather than daughter-cell separation. Fig. 4H uses initial and terminal coordinates with a common orthographic camera and scale.

### Implemented viral generation rules and host interfaces

The frozen current source specifies gene-level G + RNAP → TX and TX → G + RNAP + R + event reactions, and R + ribosome → RB and RB → R + ribosome + P + event reactions. Completion constants depend on annotated nucleotide or amino-acid length; current P1 structural weights further modify translation completion. The r12 replication reaction emits an event, and the hook constructs the daughter topology and markers. Packaging applies configured protein and topology-availability gates, consumes structural proteins and produces recorded placements (Table S6).

The source also implements post-event resource accounting: completed events request sequence-dependent nucleotide, amino-acid and energy costs. Host routines pay NTP/dNTP costs from metabolite pools and translation costs through charged-tRNA reactions. Each request is capped at 5,000 per call and applies only to an existing cost key; insufficient nucleotide pools retain unpaid balances and are set to one. These are implemented host interfaces, not direct pre-reaction substrate gates or evidence of uncapped global stoichiometric closure. Table S6 identifies exact functions and frozen source IDs, and Table S7 retains complete annotated input composition. The saved input hash identifies the base P1 gene-resolved script; Table S6 records the provenance of the other inspected source files.

### Gene states and initial spatial geometry

The eleven annotated P1 loci retain their inclusive coordinates and strand assignments. Sequence composition is reported independently of dynamic consumption. For each locus, all sampled R, RB and P states were retained, including zeros; main heatmaps use separately labelled state-specific log10(1 + count) bounds and supplementary plots show raw counts. R, RB and P peaks were calculated from cross-locus sums at the saved sample times.

Chromosome coordinates were converted using the inherited mapping x_nm = x_XYZ/10 + c, with c = (320, 320, 640) nm, and mapped to voxel indices by floor(x_nm/10). Site arrays were transformed from stored z–y–x order to x–y–z where required. Host protein abundance includes names matching P_ followed by four digits; the corresponding _TC cost states were excluded. The initial geometry is bound to the retained parent record; Fig. S1 uses the full coordinate-mapping and site-slice tables. Display cutaways and projection occlusion do not change the full object populations used for quantitative calculations.

### Conditional membrane geometry and complete bodies

The new spatial model takes the parent 1236-s site array and all eleven saved assembly events as fixed inputs. A complete body is the union of a radius-2 lattice head, an axial tail and a radius-1 baseplate, with 47 unique occupied sites. Nominal inherited geometry is a 50-nm head diameter, 80-nm tail length, 10-nm tail diameter and 20-nm baseplate diameter. These discrete dimensions define a coarse-grained tailed particle on the 10-nm grid. The early P1 isolation study reported a head diameter of approximately 28 nm [26], so the inherited 50-nm head parameter is not a reconstruction of that measurement. The 47 sites represent occupied volume rather than individual structural proteins; the packaging stoichiometry is specified independently. Each body’s identity, assembly center *c* and orientation *o* are preserved. The scheduled time *t*_s_ is supplied by the parent placement record; accepted birth time *t*_b_ is assigned by the spatial continuation when admission succeeds. Deferred admission preserves the object ID and retries without creating a new placement event.

The initial volume V comprises non-extracellular sites. Its membrane is M = V minus E26(V), where E26 denotes erosion by a 3 × 3 × 3 neighborhood. The initial reconstruction contains 9,842 membrane sites at 1236 s. At birth, the body and its one-voxel neighborhood are united with V and the membrane is recalculated. A birth that conflicts with another body or would make a new membrane intersect an existing body is delayed and retried at the next second. Re-enclosure of a released body is also rejected. The rule calculates geometry but includes no elastic-energy or lipid-supply balance.

Contact sites accumulate from the intersection of the pre-update membrane with each birth body and its six-neighbor dilation. Damage increments use the archived weights of 0.01 per new contact site and 0.0005 per retained object per second. The displacement-site, damage and capacity-failure thresholds are 160, 1.2 and 3, respectively. Contact counts are recomputed under the new definition and are not substituted for the historical marker displacement variable. Archived forced-placement events supply the capacity-failure input; the third such event triggers the gate at 1301 s. After opening, membrane sites intersecting a Euclidean dilation of the contact set are removed. The predeclared main radius is three voxels; radii two and four provide sensitivity calculations. This is a neighborhood-dilation radius, not the radius of a measured circular aperture. All membrane versions and their added/removed sites are saved.

### Stochastic translation and the release criterion

Before opening, bodies remain at their assembly sites. After opening, each active object has six axial translation channels with rate D/h^2^, where D = 10^-13^ m^2^ s^-1^ and h = 10 nm, giving 1000 attempts s^-1^ per direction. Waiting times follow the aggregate exponential proposal process and the object and direction are selected uniformly, following continuous-time stochastic simulation principles [27,28]. Rejected proposals consume time. Moves causing body overlap, intact-membrane intersection or lattice-boundary crossing are rejected; the outer lattice boundary is reflecting by rejection. No force guides motion toward a pore.

Release occurs only when every occupied body site is outside the current cell volume. Released bodies continue moving extracellularly, with re-entry proposals rejected to implement the irreversible readout. The parent releasePerSecond marker rule is not used to generate these exits. Mobility is inherited from the previous marker model and is not calibrated for the rigid body. Rotation, hydrodynamics, host crowding and feedback to the upstream reaction model are omitted.

### Sensitivity calculations and saved state verification

All nine combinations of radii 2, 3 and 4 voxels and seeds 0, 1 and 2 were calculated through 1480 s. The main display uses seed 0 and radius 3, selected before examining release outcomes. Each continuation saves 245 one-second states from 1236 through 1480 s, complete membrane versions, birth and delay events, and exact release times. Every saved state was independently checked for unique occupied sites, absence of membrane intersections, complete extracellular occupancy of released objects and assembled = retained + released. All membrane revision logs were replayed against saved arrays.

The first post-opening second of the main continuation was replayed with every accepted event retained and the original random-number sequence preserved. Collision and membrane-intersection checks, endpoint coordinates and release times were verified against the saved continuation. This replay supplies the three states in Fig. 4D.

### Visualization and reproducibility

Figures use embedded Arial. Numerical analysis and plotting used NumPy, SciPy and Matplotlib; the spatial kernel used Numba [29,30,31,32]. The analysis environment used Python 3.11, NumPy 2.4.6, SciPy 1.17.1, Matplotlib 3.10.9, Numba 0.67.0, h5py 3.16.0 and Pillow 12.2.0. Scripts, input/output hashes and frame mappings are retained with the computational record [23,24].

Spatial views use orthographic cameras and identified membrane display cutaways. Fig. 4D retains a fixed view, scale and rendering settings across three event states. The rear surface shows the 118-site opening and its 38-site intact rim; the opening pointer identifies a removed site. Orange and purple mark head and tail/baseplate sites independently of inside/outside classification. Occupancy bars classify all 47 body sites against the complete three-dimensional volume. The blue path follows recorded head-anchor positions, including returns. Fig. 4B,C show sections, whereas Fig. 3C and the rule schematics show projections. Fig. 3D evaluates a one-site proposal for the complete body; Fig. 3E illustrates the same contain, shell and pores operations used by the continuation. Voxel glyphs represent coarse-grained lattice occupancy. Video S1 retains its P1 source-time mapping; Video S2 uses all 1237 historical L1 snapshots through 4944 s. Molecular glyphs denote saved positions rather than persistent molecular identities. Display settings and exact frame mappings are retained in the computational record.

### Process accounting and object timing

P1 protein balances use all recorded translation multiplicities, the consumed_structural_subunits field in every packaging entry, and the sum of V_P1_P_* entries in each new_by_species removal dictionary. Cumulative terms at time t include events with time_s ≤ t. Their difference is compared with the saved aggregate P inventory at every checkpoint. Separate mRNA binding, completion and degradation fluxes remain incompletely resolved; they are not inferred from the protein balance. The pre-opening permeability log explicitly excludes viral species, whereas the post-opening log records their removal. Packaging entries and physical placements represent corresponding operations and are not added as independent synthesis counts.

For each spatial object, accepted birth time *t*_b_ is read from birth_events.json and states.npz. With opening time *t*_open_ = 1301 s, we calculate *T*_total_ = *t*_exit_ − *t*_b_, *T*_wait_ = max(0, *t*_open_ − *t*_b_), and *T*_post_ = *t*_exit_ − max(*t*_b_, *t*_open_). Their additive identity is checked object by object. Exact event times are used; missing exits remain censored. All nine conditions and eleven identities per condition are retained (Fig. S7).

For the paired exit-readout analysis, the anchor is the recorded head/assembly position *c*, an occupied zero-offset body site. First anchor crossing occurs when *c* lies outside the current volume; complete-body exit occurs when all 47 sites lie outside. Their time difference and intervening anchor returns are recorded. Each condition is replayed from its original seed and saved admission/boundary schedule, preserving random draws and verifying checkpoint states, proposal counts and exit times. All 99 condition–object records are included, with no censored exits. Cumulative first-crossing counts are absorbing readouts; instantaneous anchor-outside counts allow returns. Timing is also measured from max(*t*_b_, *t*_open_), with undefined ratios retained as missing. This comparison changes the readout definition on unchanged full-body dynamics (Table S8).

### Numerical reference tests

The frozen spatial translation kernel is tested independently of the production geometry. Reaction–diffusion results can depend on discretization [33,34,35], so these tests address specific movement rules rather than convergence of the full host model. Free single-site walkers use a 25^3^ lattice, a central starting site, rate 2 s^-1^ per direction and duration 0.2 s. All seeds 0– 4095 are retained. The analytical mean-squared displacement is 6kt = 2.4 site^2^; a six-standard-error acceptance band was specified before execution. Both a single 0.2-s interval and two 0.1-s intervals are tested, with no boundary rejections. Additional tests retain the full 47-site geometry within a closed cubic boundary and then remove that boundary, using all seeds 0–31 in each case. Occupancy, release classification and proposal accounting are checked independently. These benchmarks test reference translations and interval scheduling (Table S5).

The finite-opening reference uses a 19 × 19 × 45 lattice, a one-site planar wall at z = 23, a fixed axial 47-site body and opening radii of 0, 1, 2 and 3 lattice sites. An independent configuration-space construction marks forbidden body centers by subtracting every body offset from every obstacle, then performs exhaustive six-neighbor breadth-first search. The first two openings are inaccessible and the latter two have explicit legal path certificates. The production movement predicate agrees with this independent construction. All seeds 0–31 are additionally run for 20 s per opening at 80 attempts s^-1^ per direction, producing 0, 0, 6 and 17 exits, respectively. Seeds and outcomes are not filtered. Accessibility is established by the independent search, not by observing a stochastic exit or selecting an empirical pore threshold. These tests validate the movement predicate for the defined lattice geometries (Table S5).

### Quantification and statistical analysis

P1 and L1 each contribute one archived upstream trajectory under different configurations (Table S4). The nine spatial conditions share the P1 upstream events. The 99 paired readouts comprise eleven object identities in each of nine conditions; pooled summaries describe this set rather than independent production outcomes. Biological-model trajectories are reported descriptively, without inferential tests, P values or confidence intervals. Standard errors in the free-diffusion benchmark quantify Monte Carlo sampling of a known numerical reference. Medians and interquartile summaries describe the indicated objects within a frame or condition; parameter comparisons retain individual trajectories.

Radial distributions use all initial objects relative to the fixed model center and are not shell-volume-normalized densities. Genome dispersion is the root-mean-square distance from each checkpoint’s genome-marker centroid. Retained-to-membrane distances are unsigned Euclidean distances to the nearest membrane-site coordinate. Packaged-to-retained distances use directional nearest-neighbor matching without a one-to-one constraint. Distances are undefined when the required source or target population is absent; missing values remain NA and are not bridged across missing checkpoints. Time points, genes, objects and log entries are not treated as independent simulation replicates.

