## Supplemental Figures and tables for "Constructing Virtual Viruses in A Spatial Minimal Cell"

Document S1 contains Figures S1–S7, Tables S1–S9 and the legends for Videos S1 and S2. Figures S1–S3 and Tables S1–S3 use the P1 parent record r12ce_g1_2500s or its conditional continuations. Figure S5 retains the parent spatial maps and all descriptive distance analyses associated with Figure 3. Figure S4 and Table S4 separately document one historical L1 configuration. The nine membrane continuations share the P1 parent record. These materials report computational modeling and visualization only, with no wet-lab experiments or human-related data.

#### SUPPLEMENTAL FIGURE TITLES AND LEGENDS

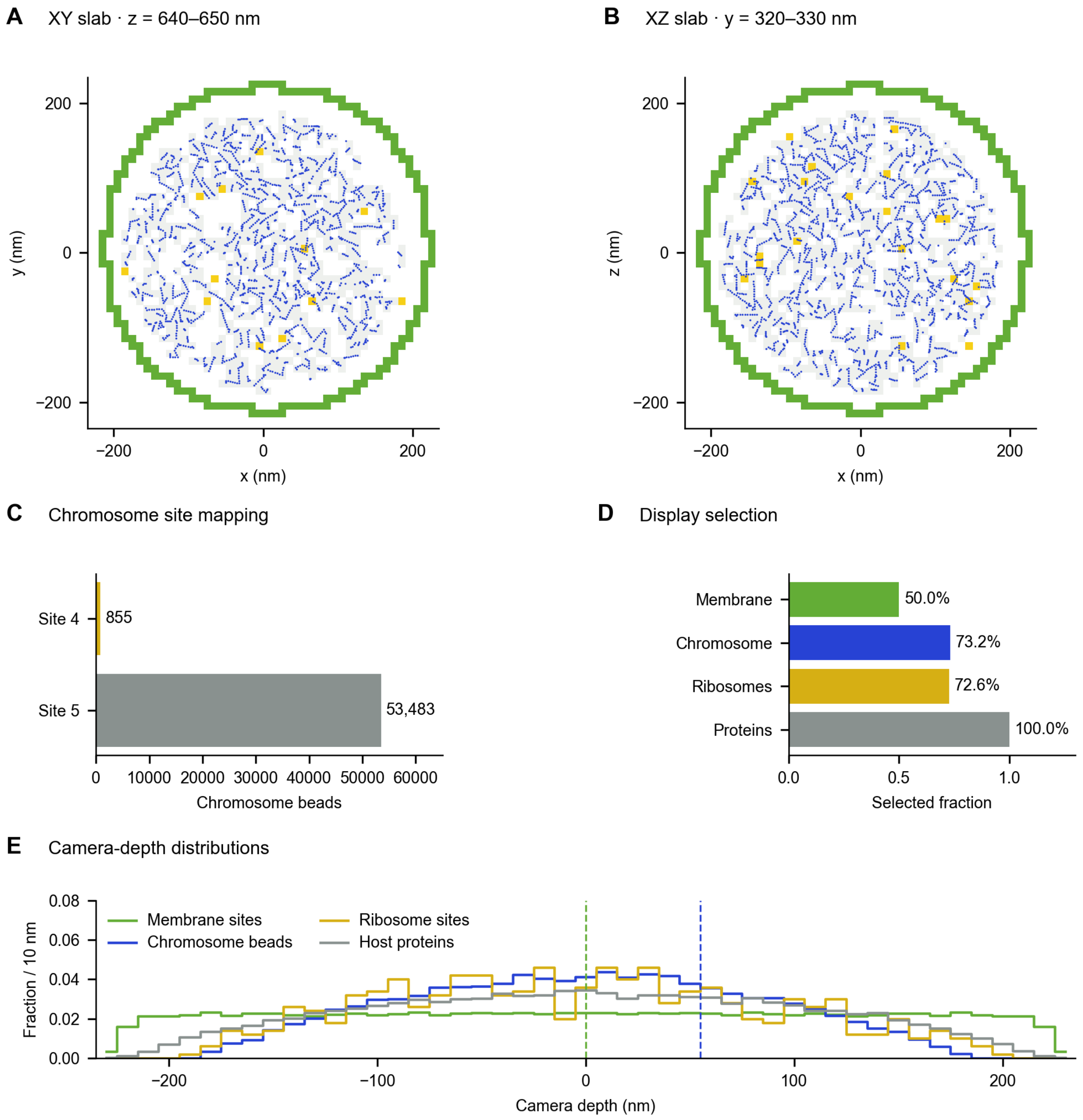

**Figure S1. Coordinate mapping and display selection for the current initial geometry Related to Figure 1**

(A,B) Orthogonal lattice sections with chromosome beads in the corresponding 10-nm slabs: z = 640–650 nm (A; 2,141 beads) and y = 320–330 nm (B; 2,298 beads). Colors distinguish the inherited site classes. The coordinate conversion and lattice indexing are those used for Fig. 1. (C) Site assignments for all 54,338 chromosome beads: 855 map to site type 4 and 53,483 to type 5, with none in other classes. The two nonzero classes are displayed; the full source table also verifies zero beads in other classes. This is an internal coordinate-consistency check; no independent registration ground truth is available. (D) Full and display-selected object counts. The cutaway retains 4,513 of 9,026 membrane sites (50.0%), 39,749 of 54,338 chromosome beads (73.15%), 363 of 500 ribosome-center sites (72.6%), and all 65,083 host protein particles. Display selection is not a count estimate and does not guarantee that every selected object is visible after projection. (E) Camera-depth distributions of all membrane sites, chromosome beads, ribosome-center sites and host protein particles, with the membrane 0-nm and chromosome/ribosome 55-nm display thresholds indicated. Membrane selection uses the camera-facing half cut. All quantitative main-figure distributions use the full populations rather than the display-selected subset. The source geometry was re-extracted from the current parent run; historical initial coordinates from a different run were not reused. These panels describe one initial configuration without inferential statistics.

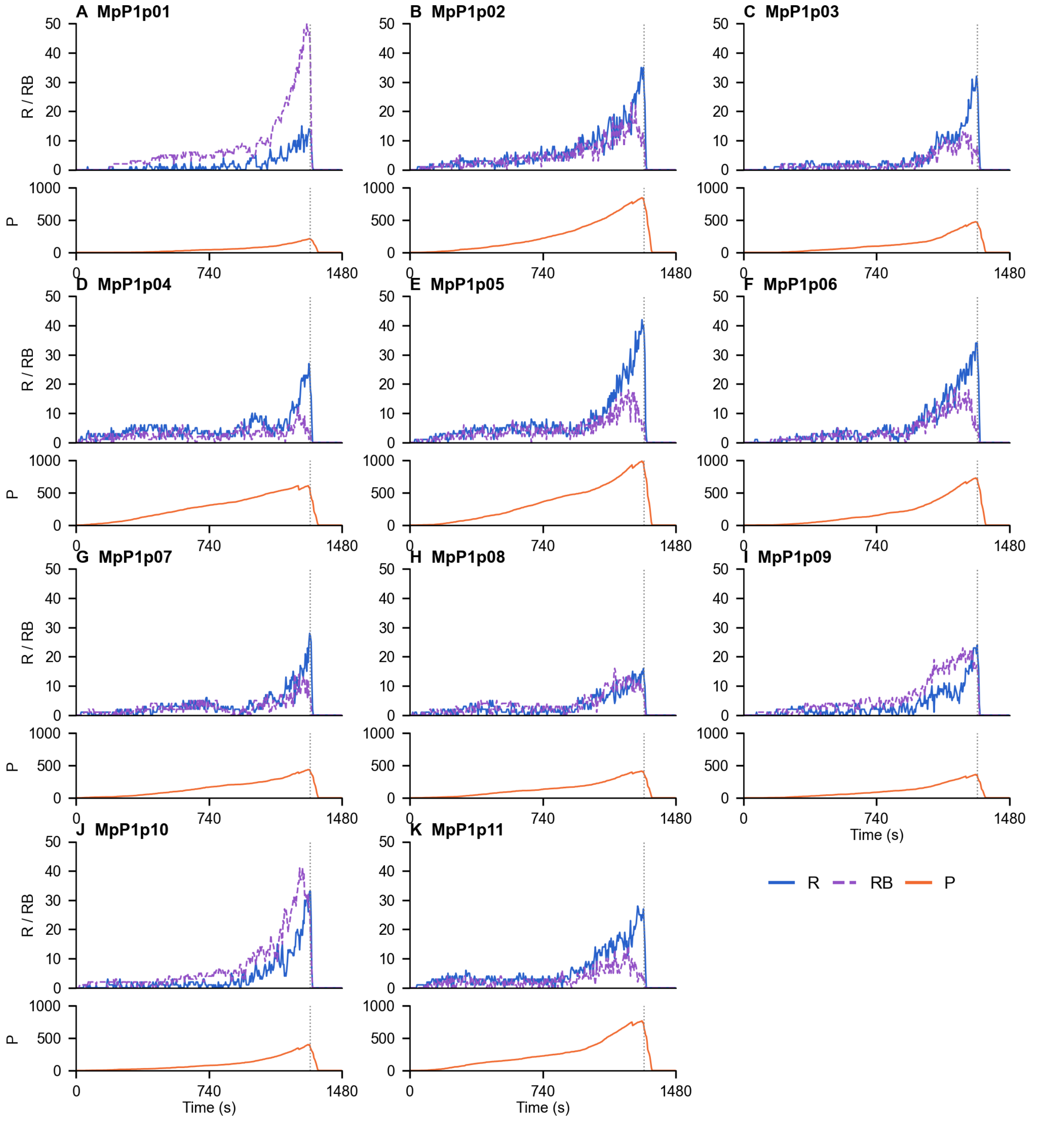


**Figure S2. Complete gene resolved state trajectories Related to Figure 2**

(A–K) Raw sampled counts for MpP1p01 through MpP1p11, respectively. Each locus has an upper axis for R (blue, solid) and RB (purple, dashed), and a lower axis for P (orange). R is the mRNA state outside the ribosome-bound complex, RB is the ribosome-bound mRNA state, and P is the protein state. All 371 saved samples at 4-s intervals from 0 to 1480 s are shown for each state and locus, totaling 12,243 observations. Upper axes share limits of 0–50 and lower axes share limits of 0–1000. Lines connect recorded samples without smoothing; terminal zeros are retained. Dotted vertical lines mark the archived opening flag at 1301 s. These are raw counts, not normalized profiles or replicate means. The independent upstream simulation count is one; no statistical tests or uncertainty intervals are included.

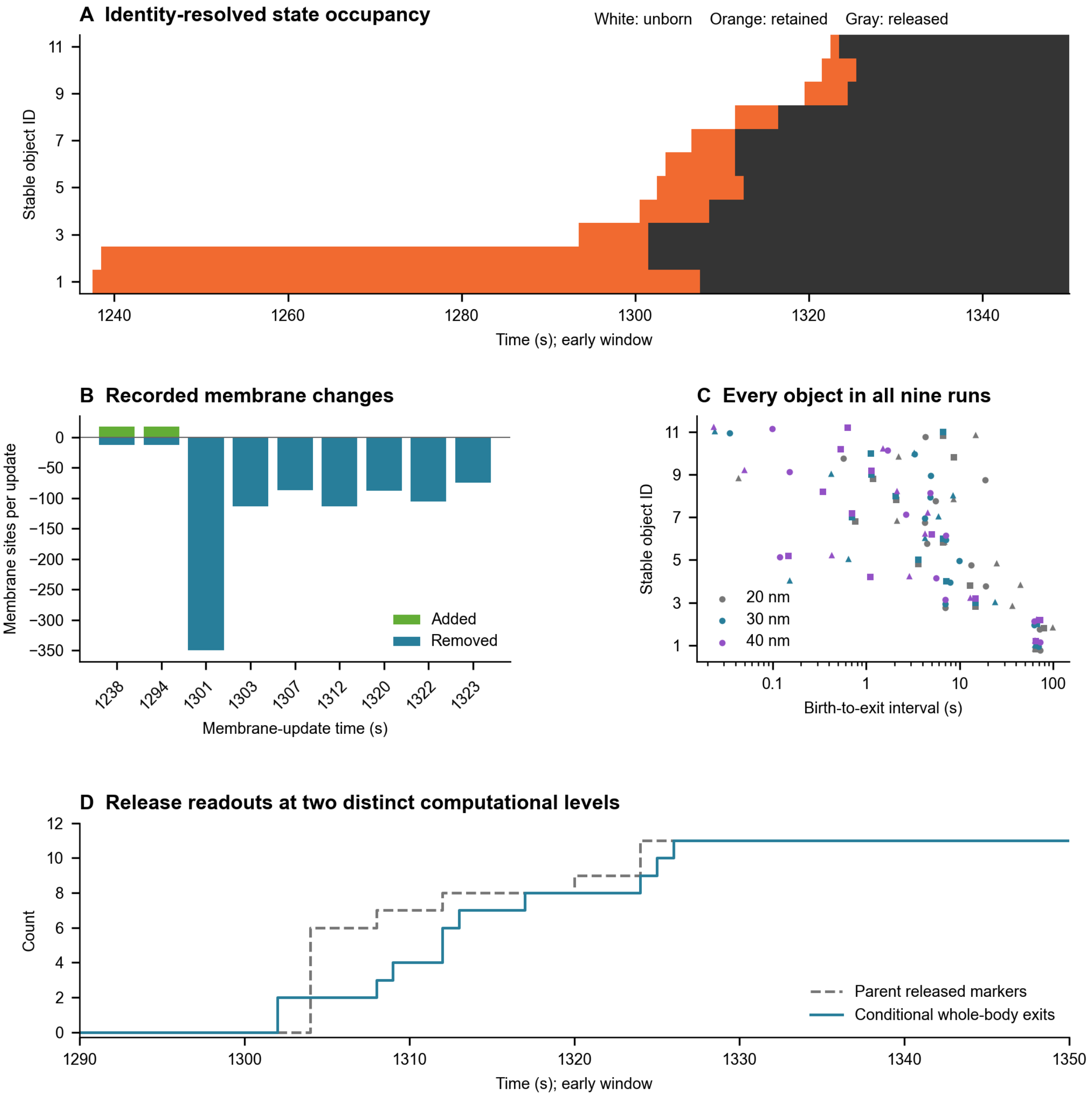


**Figure S3. Object histories and membrane sensitivity calculations Related to Figure 4**

(A) One-second state occupancy for stable object identities #1–#11 in the main continuation (30-nm contact-neighborhood radius, seed 0). White denotes not yet assembled, orange denotes retained and dark gray denotes completely released. The display enlarges 1236–1350 s; all stored states extend to 1480 s. (B) Added and removed membrane-site counts for every membrane revision after initialization in that continuation, plotted with their recorded update times. Negative bars denote removals, not negative site inventories. Revisions can combine enclosure and pore changes; cumulative pore size is reported separately in Fig. 4F. (C) Birth-to-complete-exit intervals for every object in all nine radius–seed combinations (99 points). Radius is encoded by color and seed by shape. The logarithmic time axis displays individual intervals, without aggregation or inferred uncertainty. Small deterministic vertical offsets distinguish conditions sharing an object identity. Birth time is the accepted assembly time, including any delay. (D) Cumulative parent released-marker readouts and conditional complete-body exits for the main continuation, shown over 1290–1350 s. The former are sampled every 4 s; the latter derive from computed complete-body states. Their proximity is not a calibration test or evidence that the original marker transition contained a physical path. All nine continuations use the same parent events; they are not independent whole-cell replicates.

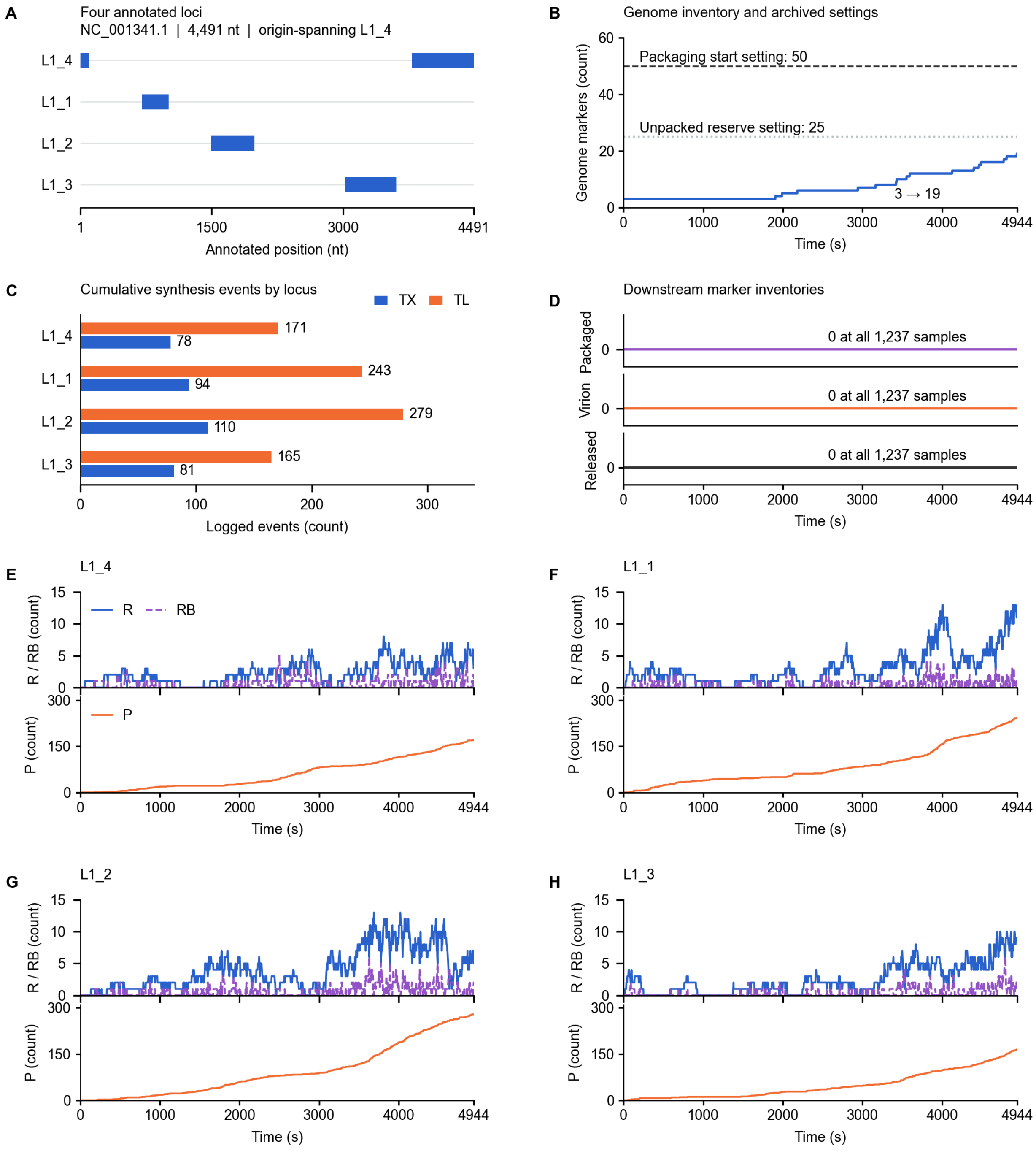

**Figure S4. Historical L1 expression and downstream state inventories Related to Figures 1 and 2**

(A) Four loci from the saved NC_001341.1 annotation on a 4491-nt circular genome, shown on a linear coordinate axis. L1_4 comprises positions 3785–4491 and 1–94; the two segments represent one origin-spanning locus. Locus order follows the archived CDS table. (B) All sampled genome-marker inventories, increasing from 3 to 19. Horizontal lines show the archived packaging-start setting of 50 and unpacked-genome reserve setting of 25; they are configuration values, not measured biological thresholds. The exact historical implementation has not been recovered. (C) Total logged transcription (TX, blue) and translation (TL, orange) events for each locus, summed over 1043 event-log rows. Totals are TX = 363 and TL = 858; bars represent one record, not means. (D) Packaged-genome, virion and released marker inventories on separate aligned axes. All three rows are present and zero at all 1237 samples; separating the axes prevents coincident zero traces from hiding a state. (E–H) Complete raw R, RB and P trajectories for L1_4, L1_1, L1_2 and L1_3, respectively. R is unbound mRNA, RB is ribosome-bound mRNA, and P is protein. Upper axes show R (blue solid) and RB (purple dashed); lower axes show P (orange). Upper limits are shared across loci (0–15 for R/RB; 0–310 for P). Lines connect saved samples without smoothing. All 1237 samples from 0 to 4944 s are retained; the requested 7200-s duration was not fully observed. Event totals use summed multiplicities, as for P1. The state panels use raw inventories with explicit axes; the independent P1 and L1 ranges are not normalized across loci. Each P trajectory equals its cumulative logged translation at the sampled times; the genome inventory equals 3 plus cumulative logged replication (16 events). These panels describe one L1 trajectory under its saved configuration (Table S4).

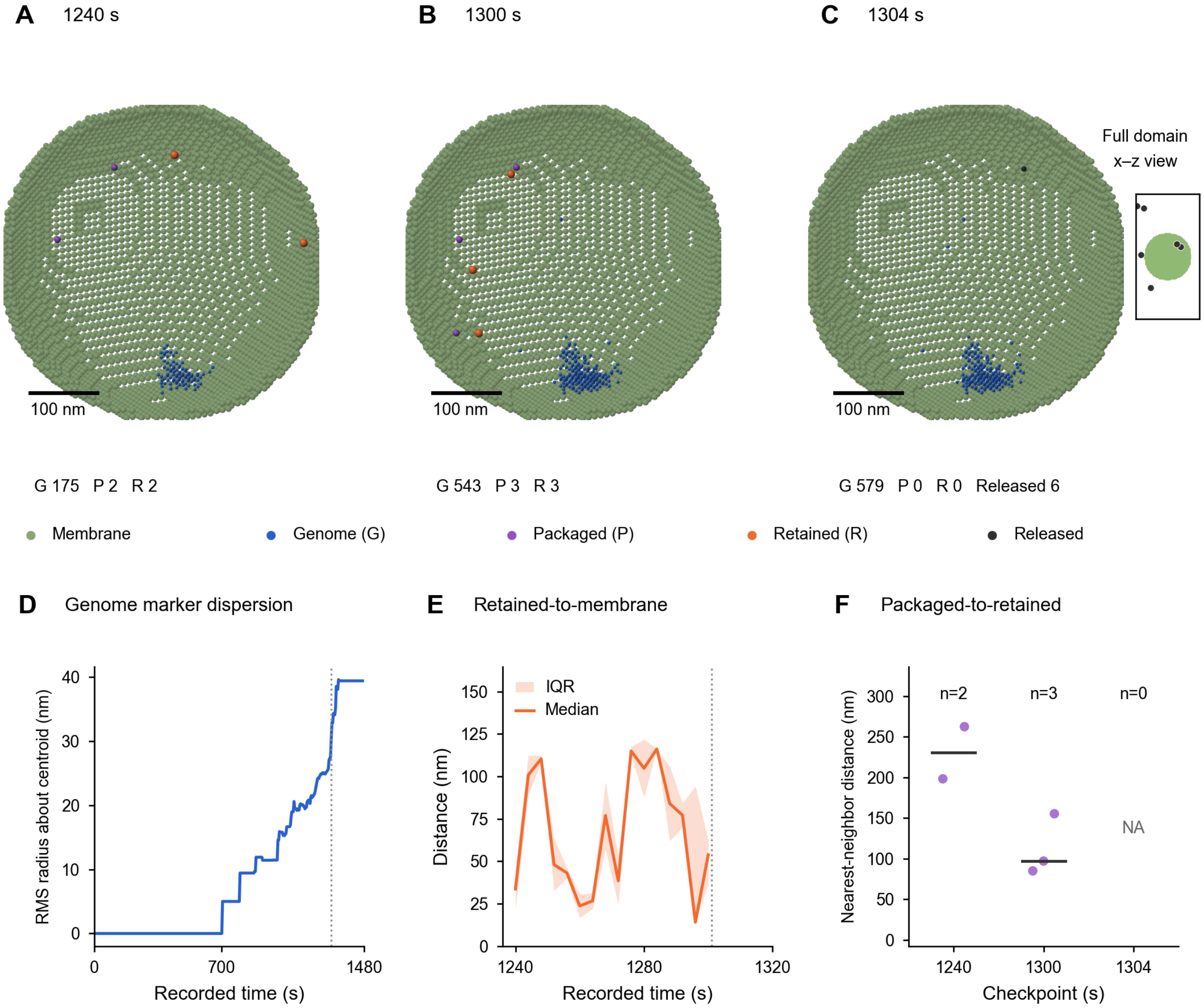


**Figure S5. Spatial marker distributions and complete distance records Related to Figure 3**

(A–C) Spatial records at the first checkpoint containing packaged and retained markers (1240 s; A), the last stored checkpoint preceding the opening flag (1300 s; B), and the first checkpoint with nonzero released readout (1304 s; C). The three cell views share the same orthographic projection, spatial window, displayed scale, glyph radii, and membrane cutaway. Genome (blue), packaged-genome (purple), and retained (orange) counts are 175/2/2, 543/3/3, and 579/0/0, respectively. Six released markers are recorded at 1304 s. The full-domain x–z inset in C includes all six released coordinates, including those outside the cell-view window; the inset has a separate scale and is not a trajectory plot. Counts refer to complete marker populations, including glyphs occluded in projection. Green glyphs indicate contemporaneous membrane sites; initial host chromosome coordinates are not inserted into these later frames. Cell-view scale bars, 100 nm.

(D) Root-mean-square distance of genome markers from their contemporaneous centroid across all 371 stored checkpoints (0–1480 s, 4-s intervals). Each marker contributes equally. A single marker gives a radius of zero by definition; the final value is 39.41 nm. This is the spatial dispersion of discrete genome-state markers, not the radius of gyration of a resolved chromosome polymer. (E) Median and interquartile range (shading, 25th–75th percentiles) of the unsigned distance from each retained marker to its nearest contemporaneous membrane-site coordinate. The metric is defined at 16 stored checkpoints and is undefined at the other 355 checkpoints, which contain no retained marker. The focused time window contains all defined samples; missing values are not replaced with zero or bridged across missing samples. The band describes within-checkpoint marker heterogeneity, not uncertainty across independent runs. (F) Distances from each packaged marker to its nearest retained marker at the three displayed checkpoints. All packaged markers are shown (n = 2, 3, and 0); horizontal lines indicate medians of 230.37 and 96.95 nm at the first two checkpoints. NA denotes undefined distances at the third checkpoint, where both marker classes are absent. Horizontal point offsets are for visibility only. Nearest-neighbor matching is directional and not one-to-one; multiple packaged markers may share a nearest retained marker. These distributions do not track the same physical particle through time.

All panels describe one P1 trajectory. Vertical dotted lines indicate opening at 1301 s; lines connect recorded summaries without smoothing. Changes in D–F reflect both marker positions and population composition. Distances use nominal 10-nm lattice coordinates and the complete marker populations, independently of display cutaways. Source data comprise all checkpoint summaries, individual retained-to-membrane distances, selected marker coordinates including all released markers, and individual cross-state distances.

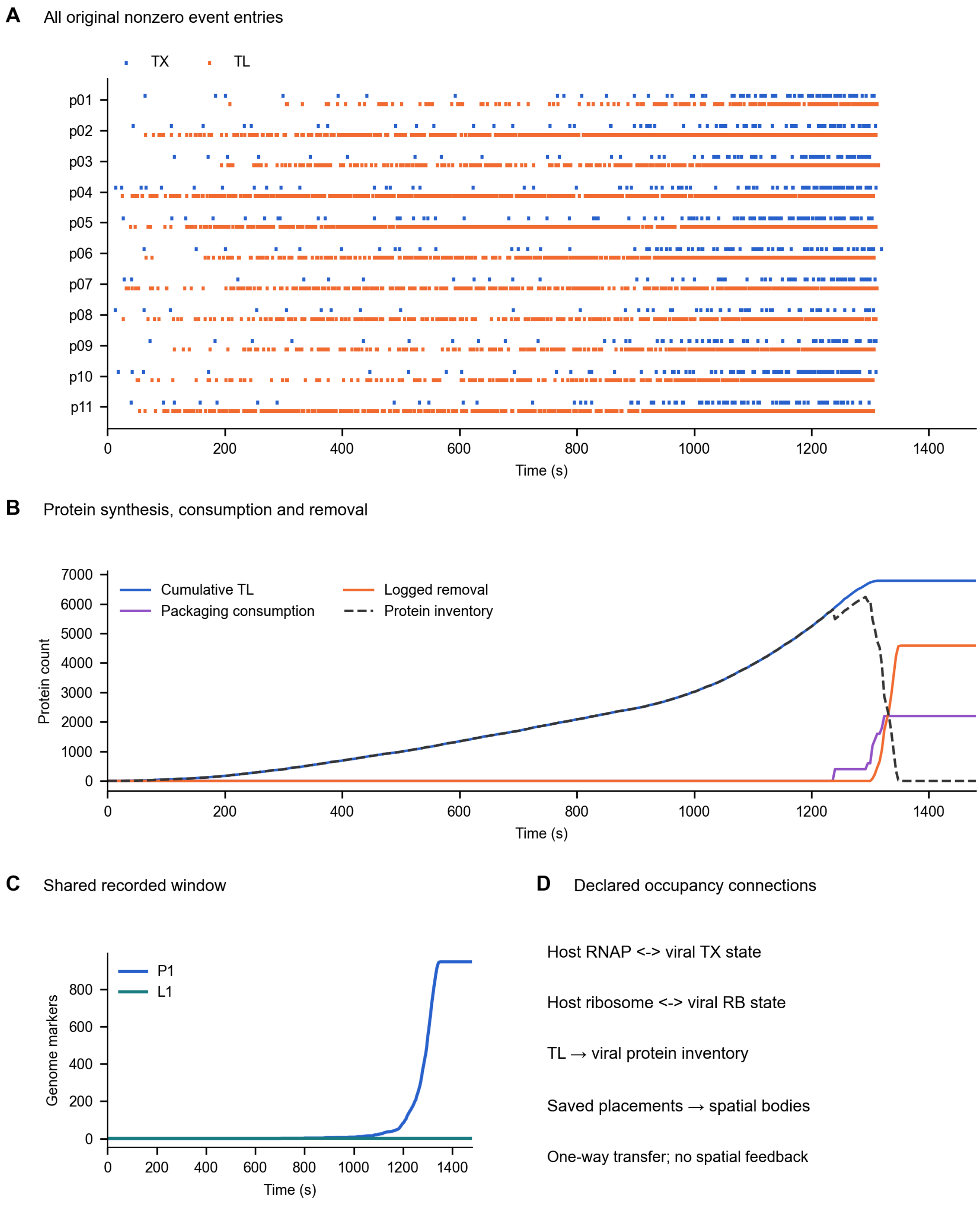


**Figure S6. Expression events, protein fates and host interfaces Related to Figures 1 and 2**

(A) All 5,450 nonzero P1 TX/TL log entries plotted at recorded times. Marks denote entries; weighted event multiplicities are analyzed in Fig. 2C,D. (B) Cumulative translation, configured packaging consumption, logged removal and instantaneous protein inventory across all 371 checkpoints. Translation minus the two losses reproduces inventory exactly. (C) P1 and L1 genome inventories within their common 0–1480-s recorded window, without matched-condition inference. (D) Configuration-declared and current-code-supported RNAP/ribosome occupancy connections, and the one-way transfer of saved placements into the spatial layer. Current source is not substituted for an unbound historical executable. No energy-consumption or bidirectional-feedback arrow is implied.

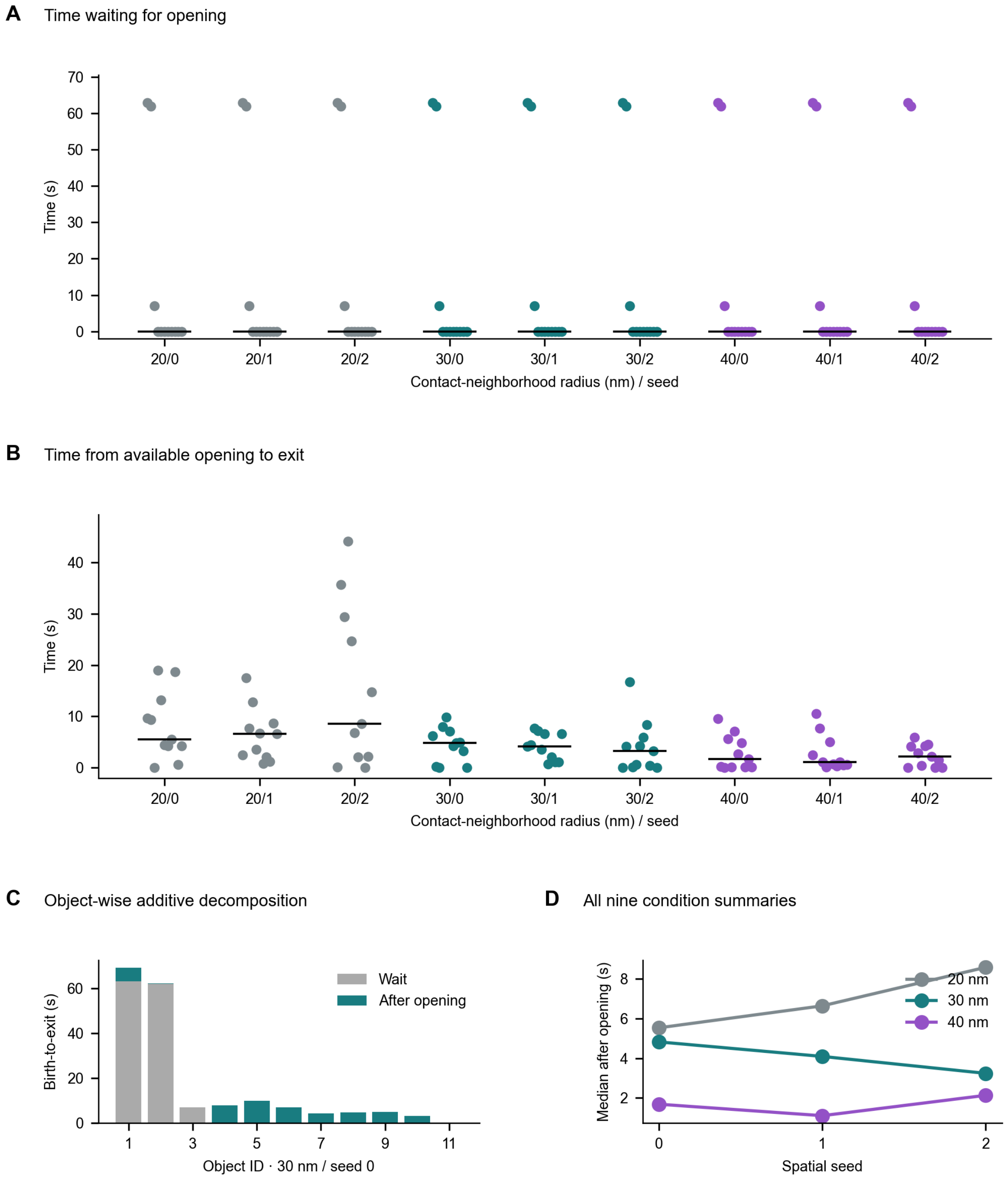


**Figure S7. Accepted-birth and opening-dependent release intervals Related to Figure 4**

(A,B) Waiting and post-opening intervals for all eleven objects in each of the nine conditions. Each dot is a stable object identity; deterministic horizontal offsets separate marks. Black segments are within-condition medians, not confidence intervals. (C) Additive waiting and post-opening times for the main 30-nm, seed-0 condition. (D) Median post-opening intervals for all conditions. Object birth uses actual acceptance, including the delayed event; none of the 99 histories is censored within the saved window. Condition medians need not add to a median total. These objects and conditional runs do not supply independent upstream replicates.

### SUPPLEMENTAL TABLES

#### *Table S1. Parent record events and accounting checks Related to Figures 1 and 2*

| Readout | Recorded value | Interpretation |
|---|---|---|
| Saved states | 371; 0–1480 s every 4 s | One parent trajectory |
| Requested duration and stored stop | 2500 s; 1481 s | Stop follows configured 180-s post-opening window |
| First TX / TL events | 13 / 25 s | Logged event times |
| First replication event | 703 s | Topology replication |
| Packaging / physical placements | 11 / 11; 1238–1323 s | Corresponding operations, not 22 particles |
| Nonzero TX/TL log entries | 5,450 | Entries can contain multiple events |
| Cumulative TX / TL events | 858 / 6,785 | Sum of recorded event values |
| Cumulative replication events | 957 | Distinct from endpoint genome inventory |
| Opening flag | 1301 s | Configured diagnostic gate |
| Endpoint genome / packaged / retained / released | 947 / 0 / 0 / 11 | Marker inventories at 1480 s |
| Instantaneous sampled packaged / retained peak | 3 / 3 | Not cumulative production |
| Accounting identities | 3 × 371 = 1,113 checks; zero mismatches | Genome and particle marker balances |

The identities are 1 + cumulative replication − cumulative packaging = genome; cumulative physical placement = retained + released; and cumulative packaging = packaged + released. They do not constitute a balance for all cellular molecules. The first nonzero released sample is at 1304 s and the preceding sample at 1300 s is zero; these samples do not specify an exact original release-event time.

#### *Table S2. Model settings and scope Related to Figures 1 to 4*

| Setting | Value | Computational role |
|---|---|---|
| Genome input | NC_002515.1; 11,660 bp; 11 loci | Annotation-defined P1 program |
| Initial genomes | 1 | Post-entry initial condition |
| placementSeed / dnaRngSeed | 11012 / 42 | Saved parent settings |
| assemblyRateOverride | 0.02 | Saved parameter, not calibrated kinetics |
| structuralSubunitTotal | 200 | Topology-packaging gate |
| Packaging reserve / fraction beyond reserve | 150 molecules / 0.03 | Saved topology-packaging settings |
| Physical-particle target | 20 | Configured target; observed total is 11 |
| Parent releasePerSecond | 4 | Marker conversion rule only |
| Post-opening observation | 180 s | Parent stop setting |
| Conditional initial lattice | 1236 s; 9,842 membrane sites | Fixed input to continuation |
| Lattice spacing / complete body | 10 nm / 47 occupied sites | Rigid-body representation |
| Nominal head / tail / tail width / baseplate | 50 / 80 / 10 / 20 nm | Inherited assumed geometry |
| Contact-site / damage / capacity gates | 160 / 1.2 / 3 | Phenomenological opening inputs |
| Damage increments | 0.01 per contact; 0.0005 per retained body per second | Recomputed contact definition |
| Contact-neighborhood radii | 20, 30, 40 nm | Dilation radius, not aperture measurement |
| Spatial seeds | 0, 1, 2 | Main continuation uses radius 30 nm and seed 0 |
| Diffusion coefficient | $10^{-13}$ $m^2$ $s^{-1}$ | Inherited marker mobility, uncalibrated for body |

| Setting | Value | Computational role |
|---|---|---|
| Release criterion | All 47 sites outside current volume | Irreversible readout; re-entry rejected |
| Motion and coupling | Translation only; no parent feedback | Rotation, mechanics and host crowding omitted |

The full archived configuration accompanies the source data and takes precedence over this selected-parameter table. Capacity-failure inputs use archived forced-placement events. Re-enclosure of released objects, body overlaps and membrane intersections are rejected. Nine conditional calculations do not replace independent whole-cell simulations.

***Table S3. All conditional spatial continuations Related to Figure 4***

| Radius (nm) / seed | Final released | First exit (s) | Last exit (s) | Peak fully inside | Delays | Pore sites |
|---|---|---|---|---|---|---|
| 20 / 0 | 11 | 1301.032 | 1338.717 | 6 | 0 | 541 |
| 20 / 1 | 11 | 1303.468 | 1330.658 | 5 | 0 | 541 |
| 20 / 2 | 11 | 1301.075 | 1345.070 | 6 | 1 | 541 |
| 30 / 0 | 11 | 1301.032 | 1325.278 | 5 | 0 | 950 |
| 30 / 1 | 11 | 1305.093 | 1329.629 | 6 | 0 | 950 |
| 30 / 2 | 11 | 1301.035 | 1325.241 | 4 | 0 | 950 |
| 40 / 0 | 11 | 1301.026 | 1323.686 | 4 | 0 | 1224 |
| 40 / 1 | 11 | 1302.109 | 1323.630 | 4 | 0 | 1224 |
| 40 / 2 | 11 | 1301.022 | 1323.507 | 4 | 0 | 1224 |

Times are in seconds and rounded to three decimals here; full-precision release times for all 99 object–condition pairs are provided in source_data/all_object_exit_times.csv. Radius denotes contact-neighborhood dilation. Each condition contains three complete intracellular objects at 1294 s, releases 11 of 11 objects, and has zero violations in 245 saved-state checks. Peak fully inside is an instantaneous one-second count, distinct from retained (which also includes partly exited bodies) and cumulative assembly. Delays counts rejected birth attempts. All conditions add 126 volume sites. The opening gate is 1301 s in every condition. None of these rows is an independent upstream replicate.

***Table S4. Historical L1 record and comparability boundaries Related to Figures 4 and S4***

| Readout | P1 parent | Historical L1 |
|---|---|---|
| Archive role | Main parent trajectory | Separate descriptive configuration |
| Input annotation | NC_002515.1; 11 loci | NC_001341.1; 4 loci |
| Initial genome markers | 1 | 3 |
| Saved samples / endpoint | 371 / 1480 s | 1237 / 4944 s |
| Requested duration | 2500 s | 7200 s |
| placementSeed / dnaRngSeed | 11012 / 42 | 70701 / 42 |
| TX / TL / replication events | 858 / 6785 / 957 | 363 / 858 / 16 |
| Final genome / packaged / virion / released | 947 / 0 / 0 / 11 | 19 / 0 / 0 / 0 |
| Packaging settings | Topology-linked settings; Table S2 | Start 50; unpacked reserve 25; conversion 5 per s |
| L1 CSV–HDF5 agreement | Not an L1 comparison | 44,532 entries; zero mismatches |
| L1 event–inventory agreement | Not an L1 comparison | 6185 checks; zero mismatches |
| Complete-body spatial continuation | Nine conditional continuations | None performed |
| Historical source identity | Executable not recovered | Recorded and current source hashes differ |

The saved L1 host geometry provides the descriptive readouts in Fig. 4H–J. All 1237 Sites checkpoints contain one connected cell domain under both 6- and 26-neighbor connectivity. The cell-domain voxel count increases from 50,397 to 95,845 on a 10-nm lattice, corresponding to 0.050397–0.095845 $\mu m^3$ including membrane sites. The final axial midplane-to-lobe area ratio is 0.799183. The historical log records ten division-associated geometry updates beginning at 4288.25 s, and the terminal division_started flag is true. This documents entry into division-associated shape updating, not completed daughter-cell separation or absence of an L1 effect. Quantification uses the mask and section definitions in Methods; snapshots and geometry versions are not independent replicates.

Genome-marker abundance was also divided by cell-domain volume at each of the 1237 shared timestamps. The initial and terminal values are 59.53 and 198.24 markers $\mu m^{-3}$, respectively, giving a 3.33-fold increase: (19/0.095845)/(3/0.050397). Genome accumulation therefore exceeds the 1.90-fold increase in cell-domain volume. The denominator includes membrane sites and is not a measured cytoplasmic volume. All timestamped values are supplied in source_data/L1_morphology/L1_volume_normalized_genome_markers.csv.

Each archive represents one configuration with one retained trajectory. These columns document provenance and non-equivalence, not an estimate of a virus-specific effect. L1 downstream marker rows and assembly-event values are recorded as zero; no separate assembly log was found, and absence of that log is not itself evidence of zero physical events. The L1 genomic annotation and packaging values are taken from archived metadata. The L1 run was collected before its requested endpoint; 4944 s is the last saved observation, not proof of successful completion. Full event rows, raw state tables, annotation, archived configuration and source hashes are retained in source_data/L1. The recorded L1 script SHA-256 is a8f8ef3a3e8e319894806254d48c9e149e288c7aa26f7edd38e46abd75f784bf; the current file has a different hash. Without the archived executable, the zero downstream readouts cannot establish biological non-assembly or isolate a causal implementation defect.

***Table S5. Independent numerical reference checks Related to Figure 4***

| Reference | Expected | Observed | Interpretation |
|---|---|---|---|
| Free translation, one 0.2-s interval | MSD 2.4 $site^2$ | 2.453125; SE 0.041834; n = 4096 | Within predeclared six-SE band |
| Free translation, two 0.1-s intervals | MSD 2.4 $site^2$ | 2.375977; SE 0.037447; n = 4096 | Within predeclared six-SE band |
| Complete body, closed boundary | No exit; 47 occupied sites | All 32 seeds checked | Geometric invariant |
| Complete body, fully opened boundary | Exit permitted; no re-entry | All 32 seeds retained | Exact results in reference_benchmarks.json |
| Existing geometry/kernel tests | All tests pass | 9 tests | Scope defined in test source |
| Finite planar apertures, radii 0/1/2/3 lattice sites | Independent body-center search: inaccessible / inaccessible / accessible / accessible | 89,838 movement comparisons; 0 mismatches | Fixed axial 47-site body; full path certificates retained |
| Same finite apertures, all 32 seeds per radius | No exit when inaccessible; exit permitted when accessible | Exits: 0/32, 0/32, 6/32, 17/32 over 20 s | A finite stochastic observation is not the accessibility proof |

Monte Carlo standard errors apply only to independent numerical reference walkers. They are not uncertainty estimates for P1/L1 trajectories. No spatial lattice-resolution convergence is asserted.

***Table S6. Implemented generation rules and host interfaces Related to Figures 1–3***

| Stage | State update | Rate or gate | Host interface | Frozen source |
|---|---|---|---|---|
| P1 TX | G + RNAP → TX → G + RNAP + R + event; R decay | Binding: 10^6 × configured rate; completion max(0.005,45/nt length); decay 0.006 s−1 | RNAP occupancy; TX-triggered sequence NTP costs | S6:1493–1531; S7:324–361 |
| P1 TL | R + ribosome → RB → R + ribosome + P + event | Binding: 10^6 × configured rate; completion max(0.005,15/aa length) × structural weight | Ribosome occupancy; amino-acid and 2 × aa-length GTP cost requests | S6:1493–1531; S7:345–361 |
| P1 r12 replication | Genome + polymerase → Genome + polymerase + event; | Configured replication rate and placement feasibility | Sequence dNTP cost requests | S6:1493–1531,1693–1779 |

| Stage | State update | Rate or gate | Host interface | Frozen source |
|---|---|---|---|---|
| | hook adds daughter topology/markers | | | |
| P1 packaging / placement | Consume configured proteins; registry consumption; marker and physical-placement operations | Protein sets, topology reserve/fraction, binomial trial and fallback | 200 structural subunits per historical event; no additional paid packaging ATP stoichiometry inferred | S6:1669–1691,1953–2088; template mismatch unresolved |
| Host cost payment | Debit NTP/dNTP; charged tRNA reactions pay translation costs | Request capped at 5,000 per call; existing keys only; unpaid costs retained | Post-event accounting; insufficient nucleotide pool set to one | S7:305–308; S2:97–141; S4:731–844 |
| Current L1 | Generic G/TX/R/RB/P; annotation-dependent replication; direct genome/protein assembly | Saved L1 parameters; no separate packaged-genome species | RNAP/ribosome occupancy and cost counters | S8:240–282,360–410; not historical L1 executable |
| Spatial continuation | Placement → accepted identity/body → translation → whole-body exit | 6 × active count × per-direction rate; full-body exclusion; explicit opening gate | One-way import, no host writeback | S9:23–94; S10:14–46; S11:10–59 |

S2 is Rxns_CME.py; S4 Communicate.py; S6 the P1 r12 topology override; S7 the base P1 gene-resolved script; S8 the generic viral script; S9 run_model.py; S10 membrane_model.py; S11 walk.py. Exact snapshots, function ranges and SHA-256 values are included in the source manifest and the accompanying implementation table. The saved P1 input manifest binds S7; it does not bind the inspected historical identity of S2/S4/S6. These specifications describe the frozen current implementation; event totals remain observations of the historical archive.

***Table S7. Complete annotated sequence composition Related to Figure 2***

| Program | Locus | nt | aa | A | C | G | T |
|---|---|---|---|---|---|---|---|
| P1 | MpP1p01 | 2085 | 694 | 825 | 250 | 314 | 696 |
| P1 | MpP1p02 | 966 | 321 | 408 | 106 | 135 | 317 |
| P1 | MpP1p03 | 1173 | 390 | 473 | 129 | 166 | 405 |
| P1 | MpP1p04 | 243 | 80 | 110 | 19 | 35 | 79 |
| P1 | MpP1p05 | 249 | 82 | 96 | 20 | 36 | 97 |
| P1 | MpP1p06 | 549 | 182 | 253 | 47 | 77 | 172 |
| P1 | MpP1p07 | 2067 | 688 | 791 | 280 | 295 | 701 |
| P1 | MpP1p08 | 1200 | 399 | 475 | 160 | 179 | 386 |
| P1 | MpP1p09 | 975 | 324 | 419 | 103 | 133 | 320 |
| P1 | MpP1p10 | 1002 | 333 | 375 | 146 | 139 | 342 |
| P1 | MpP1p11 | 261 | 86 | 126 | 44 | 32 | 59 |
| L1 | L1_4 | 801 | 266 | 309 | 160 | 131 | 201 |
| L1 | L1_1 | 309 | 102 | 129 | 53 | 43 | 84 |
| L1 | L1_2 | 492 | 163 | 202 | 97 | 67 | 126 |
| L1 | L1_3 | 585 | 194 | 217 | 96 | 81 | 191 |

Values describe input sequences. Origin-spanning L1_4 is counted once. Table S9 links P1 loci to the retained product annotations, configured weights and recorded synthesis totals.

***Table S8. Paired anchor and complete-body exit readouts Related to Figure 4***

| Radius (nm) / seed | ΔT median (range), s | Objects with anchor return | Maximum cumulative count gap | Final anchor/body count |
|---|---|---|---|---|
| 20 / 0 | 0.020 (0.000628–18.471) | 8/11 | 2 | 11/11 |
| 20 / 1 | 0.027 (0.000298–0.066) | 9/11 | 1 | 11/11 |
| 20 / 2 | 0.055 (0.000635–28.106) | 10/11 | 4 | 11/11 |

| Radius (nm) / seed | ΔT median (range), s | Objects with anchor return | Maximum cumulative count gap | Final anchor/body count |
|---|---|---|---|---|
| 30 / 0 | 0.086 (0.000161–4.654) | 9/11 | 2 | 11/11 |
| 30 / 1 | 0.094 (0.000218–6.121) | 9/11 | 3 | 11/11 |
| 30 / 2 | 0.003 (0.000032–4.416) | 8/11 | 2 | 11/11 |
| 40 / 0 | 0.004 (0.000694–5.220) | 9/11 | 3 | 11/11 |
| 40 / 1 | 0.025 (0.001571–7.063) | 10/11 | 4 | 11/11 |
| 40 / 2 | 0.006 (0.001785–3.782) | 8/11 | 2 | 11/11 |

Each condition contains all eleven accepted objects from the same parent placement set. Radius is the contact-neighborhood dilation parameter. ΔT is complete-body exit time minus the first outside event of the occupied head/assembly anchor. An anchor return means an outside-to-inside transition before complete exit. The maximum cumulative count gap compares absorbing first-anchor-crossing and whole-body-exit counts at the same time; it does not represent final yield. Final counts use 1480 s. Medians and ranges describe within-condition objects. Across the full set of 99 condition–object records, the pooled median is 24.850 ms, 80 records contain an anchor return, and the maximum cumulative count gap is four. These records reuse the same eleven input identities across nine conditions and are not independent upstream replicates. Instantaneous anchor-outside counts may decrease after returns; the cumulative first-crossing count in this table cannot. No inferential tests or uncertainty estimates are used.

Exact replay matches body centers and release flags at all 2,205 condition–time checkpoints, final proposal/rejection totals and all 99 exit times. The all-object table includes scheduled and accepted birth times, opening time, both exit readouts, return counts, post-opening intervals, ratios and censor flags. The event table stores every change in external-site count or anchor status until complete exit; together with birth and end times, it specifies the piecewise-constant readout without interpolation. No object is censored.

***Table S9. P1 locus annotations, configured weights and synthesis output Related to Figure 2***

| Locus / gene | Retained product label | TL weight | Copies per packaging event | TX events | TL events |
|---|---|---|---|---|---|
| MpP1p01 / orf1 | DNA polymerase | 1 | 0 | 75 | 219 |
| MpP1p02 / orf2 | P38 | 3 | 19 | 83 | 901 |
| MpP1p03 / orf3 | P46 | 4 | 13 | 71 | 520 |
| MpP1p04 / orf4 | P10 | 1 | 34 | 85 | 689 |
| MpP1p05 / orf5 | P10.2 | 1 | 30 | 100 | 1070 |
| MpP1p06 / orf6 | P23 | 1.8 | 19 | 84 | 780 |
| MpP1p07 / orf7 | P81 | 6 | 11 | 62 | 477 |
| MpP1p08 / orf8 | P46 | 2.5 | 14 | 52 | 449 |
| MpP1p09 / orf9 | P38.4 | 1 | 15 | 71 | 399 |
| MpP1p10 / orf10 | P38.8 | 1 | 14 | 94 | 443 |
| MpP1p11 / orf11 | P10.1 | 1.3 | 31 | 81 | 838 |

Locus-to-gene correspondence is taken directly from the retained NC_002515.1 annotation. DNA polymerase is the annotation for MpP1p01; the P-number labels for the other loci do not by themselves identify a biological role. TL weights and packaging-copy requirements come from the saved P1 configuration. Unlisted translation weights default to 1; MpP1p01 is excluded from structural consumption. The requirements sum to 200 copies per packaging event. Weights modify the length-dependent completion rate in the inspected implementation (Table S6), not the measured abundance of a natural viral protein. TX and TL entries are summed event multiplicities from the retained 0–1480-s trajectory; they sum to 858 and 6,785. The table does not assign early/late regulatory classes or estimate translation efficiency.

### SUPPLEMENTAL VIDEO LEGENDS

***Video S1. P1 replication, conditional assembly and release Related to Figures 1 to 4***

The annotated P1 movie displays genome-state accumulation, structural-protein states, conditional assembly and complete-body exit. Blue denotes genome-state markers and gold denotes structural-protein markers. The molecular segments use the retained P1 parent record; conditional assembly uses d12_s0 and release uses the main r3_s0 continuation, following the persistent identity of object #3. The established shot sequence and frame-to-source mapping are retained, with updated materials, background prominence and sparse Arial annotations. During replication, the genome-state count and parent model time refer to the mapped saved state; enlarged genome glyphs aid visibility and do not represent DNA dimensions. Free-protein glyph brightness is reduced to 28% during 11–13 s for visual emphasis; it does not encode protein consumption. These segments connect distinct computational layers through editing; they do not trace continuous molecular transport or establish a continuously coupled or biologically validated trajectory.

The designated file is P1_Annotated_65_1080p.mp4: 44 s, 1320 frames, 1920 × 1080 pixels, 30 frames $s^{-1}$, without audio. A text-free companion and a 720p review copy accompany the video package. Earlier layer-60 movie files are preserved as the prior version. Source mappings, rendering settings and file hashes are retained in the layer-65 video package.

***Video S2. Historical L1 expression and genome accumulation Related to Figures 1 and 2***

The annotated L1 movie displays all 1237 recorded spatial snapshots at 4-s intervals over 0–4944 s. Genome markers increase from 3 to 19. Blue denotes genome-state markers and gold denotes protein markers. Molecular glyphs show genome, four protein classes and their R/RB states; template, TX and event-counter states are not added as extra molecular particles. The membrane follows its sixteen recorded versions. Packaged, virion and released inventories remain zero, with no recorded opening or damage; no membrane-opening or particle-release sequence is synthesized. Snapshot coordinates are not interpolated, molecules are not linked across frames by inferred identities, and display radii, including the 7-nm genome-glyph radius, are visual symbols rather than measured molecular dimensions.

The designated file is L1_Annotated_65_1080p.mp4: 1357 frames at 30 frames $s^{-1}$ (45.233 s), 1920 × 1080 pixels, without audio. Each saved snapshot appears once in chronological order, with 60 additional hold frames at each endpoint. A text-free companion and a 720p review copy are provided. The source MinCell.lm archive is identified by its SHA-256 and retained separately; extracted caches, state tables, frame mappings and audits accompany the video package. The movie represents one historical L1 configuration; it is not a matched control for Video S1 and does not extend the observation window to the requested 7200 s.

### UPSTREAM NUMERICAL INTERVENTIONS

The upstream record contains 3,606 negative-free-count initializations clamped to zero from 1238 to 1480 s, affecting 19 host proteins and 29 tRNAs. The cumulative safe-minus-raw initialization difference is 9,967 count units, not a net molecular increment. For 3,592 records, the difference equals the same-species removal logged at the same second; fourteen exceptions are retained. Current-function replay reproduces all saved safe counts. These species are reactants in the inspected host tRNA-charging network, so setting their initial free counts to zero can change CME propensities. Complete hidden-state and reaction-availability histories are not recoverable from these logs alone.

All eleven topology-consumption records report local template lookup failures, totaling 63 among 121 registered locus slots. The recorded consumed-template field counts registry entries rather than successful RDME deletions. In the inspected implementation, templates can move or enter TX, whereas packaging searches their registered coordinates for G. Extracted-function tests reproduce a moved template surviving registry consumption and a retained TX state whose completion can regenerate G. The exact historical cause of each lookup failure remains unresolved because the r12 override and full host dependency set are not historically hash-bound. Standalone repair candidates preserve hidden pools, require stable template ownership and defer packaging while TX is active; they have not been integrated into a new production trajectory. Consequently, sensitivity of expression, replication, packaging and the eleven placement inputs to corrected synchronization and template consumption remains unquantified. Source-bound tests, all affected species, per-event differences and candidate scope are supplied with the computational records.

#### *Current-source L1 feasibility case*

A separately frozen current-source L1 case completed a prespecified 60 s, with initialGenomes = 3, placementSeed = 70701 and dnaRngSeed = 42. The 16 saved checkpoints cover 0–60 s; CSV/HDF5 comparisons for all 35 viral states yield 560 agreements and zero mismatches. All initialization attempts and the file-bootstrap-only repair are retained. The current schema omits a separate packaged-genome state; its absence is not a zero observation. This short feasibility case does not regenerate or replace the historical L1 archive and does not evaluate its downstream outcome. Configuration, event counts, state tables, source hashes and the environment record are provided in source_data/Validation63/L1_current_60s.

### DATA AND CODE MANIFEST

The source data include the figure tables, full P1 and L1 state and event records, input annotations and configurations, the nine spatial continuations, membrane versions and exact object exit times. The added L1 volume-normalized genome-marker series and the P1 annotation–weight–output table are supplied with their calculation script. Numerical reference results and implementation checks accompany the analysis code. Videos S1 and S2 and their frame-to-source mappings remain in the separate layer-65 package. File hashes and locations are listed in the delivery manifest; raw parent records are retained in the upstream archive. Selected reference data and research code are publicly available at https://github.com/dmaskhhh/VirtualVirus. Project-authored software is provided under the repository MIT license; third-party materials retain their own terms. The complete historical archives and full figure/video production records remain locally retained and are not included in that public release.
